# Inter-cellular mRNA Transfer Alters Human Pluripotent Stem Cell State

**DOI:** 10.1101/2024.06.27.600209

**Authors:** Yosuke Yoneyama, Ran-Ran Zhang, Masaki Kimura, Yuqi Cai, Mike Adam, Sreeja Parameswaran, Hideki Masaki, Naoaki Mizuno, Joydeep Bhadury, So Maezawa, Hiroshi Ochiai, Hiromitsu Nakauchi, S. Steven Potter, Matthew T. Weirauch, Takanori Takebe

## Abstract

Inter-cellular transmission of mRNA is being explored in mammalian species using immortal cell lines (1–3). Here, we uncover an inter-cellular mRNA transfer phenomenon that allows for the adaptation and reprogramming of human primed pluripotent stem cells (hPSCs). This process is induced by the direct cell contact-mediated coculture with mouse embryonic stem cells (mESCs) under the condition impermissible for human primed PSC culture. Mouse-derived mRNA contents are transmitted into adapted hPSCs only in the coculture. Transfer-specific mRNA analysis show the enrichment for divergent biological pathways involving transcription/translational machinery and stress-coping mechanisms, wherein such transfer is diminished when direct cell contacts are lost. After 5 days of mESC culture, surface marker analysis, and global gene profiling confirmed that mRNA transfer-prone hPSC efficiently gains a naïve-like state. Furthermore, transfer-specific knockdown experiments targeting mouse-specific transcription factor-coding mRNAs in hPSC show that mouse-derived *Tfcp2l1*, *Tfap2c,* and *Klf4* are indispensable for human naïve-like conversion. Thus, inter-species mRNA transfer triggers cellular reprogramming in mammalian cells. Our results support that episodic mRNA transfer can occur in cell cooperative and competitive processes(4), which provides a fresh perspective on understanding the roles of mRNA mobility for intra- and inter-species cellular communications.

---

In mammalians, there is an increasing evidence of transient mRNA transfers in divergent biological contexts (1, 5–12), particularly when cells are placed under extensive stressors such as oxidative stress and nutrient deprivation with the use of immortal cell lines (1–3). Yet, it remains unknown whether the presence and contribution of inter-cellular mRNA transfer in physiologically relevant cells such as pluripotent stem cells. Here, by employing an inter-species experimental coculture model, we tested the nature and role of mRNA exchange between human pluripotent stem cells (hPSCs) and mouse embryonic stem cells (mESCs).

Unlike conventional hPSCs including induced pluripotent stem cells (hiPSCs) and embryonic stem cells (hESCs), mESCs are maintained by the leukemia inhibitory factor (LIF) signaling activation and the inhibition of GSK3 and MEK kinases, referred to as 2i+LIF condition. Under 2i+LIF condition, primed hPSCs (H9 hESCs) alone culture lost the expression of pluripotency transcription factors Oct4 and Nanog (**Fig. S1A–D**) and failed to maintain their growth capacity (**Fig. S1E**). However, we found that when cocultured with naïve mESCs, primed hESCs continued to thrive under 2i+LIF condition over five days with some human cells maintaining Oct4 and Nanog expression (**Fig. S1A–D**). mESCs-cocultured primed hESCs then expanded after human cell sorting and passaging (**Fig. S1F**). To explore the possibility of inter-species mRNA transfer during cocultured hESC adaptation in 2i+LIF condition, we first investigated the presence of the mouse cell-derived reporter mRNA in human cells in the coculture system comprising naïve mESCs expressing EGFP (G4-2) and primed hESCs (H9) without EGFP *in situ* by RNAscope (**Fig. 1A and Fig. S2A**). By utilizing the human-specific antibody against a nuclear protein Ku80, we could readily distinguish the acceptor hESCs from the donor mESCs. The transfer of *EGFP* mRNA was detected in hESCs in the mESC coculture. At the same time, there were significantly less *EGFP* mRNA levels detected in the separate culture condition either with primed hPSC culture medium (AK02N) or with conditioned medium (CM) from mESC coculture (**Fig. 1**). Quantification of *EGFP* mRNA spots per human cell colony revealed that an average of 1.56 *EGFP* probe-positive spots were present in a hESC colony in the coculture condition (**Fig. 1B**). The *EGFP* mRNA spots were observed in both peripheral and central regions of the colonies. To estimate the transferred amounts relative to the endogenous expression levels in mESCs, we unbiasedly processed images to count the RNAscope signal particles within the cells (**Fig. S2B**). Quantification of over 400 cells revealed that the donor mESCs and acceptor hESCs contain 22.5 ± 3.8 and 1.7 ± 1.8 spots/cell of *EGFP* mRNA, respectively, indicating approximately 7.5% of donor expressed transcripts transferred into acceptor cells.

**Fig. 1.**
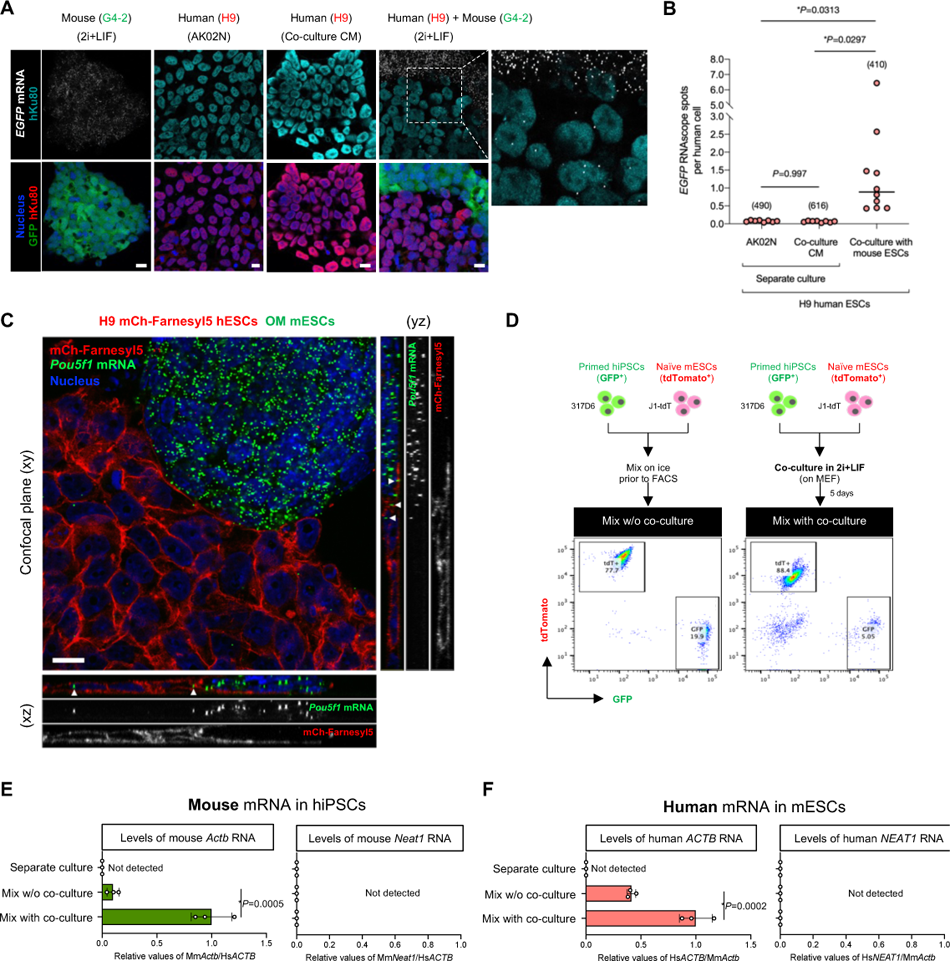
Inter-species mRNA transfer between human and mouse PSCs. (A) Representative RNAscope images of *EGFP* RNA in separate cultures of EGFP-expressing mESCs and hESCs either in primed maintenance medium or in conditioned medium of the coculture and the coculture of mouse and human ESCs. The cells were also counter-stained with anti-Ku80 antibody (human nuclei) and Hoechst 33342 (all nuclei). Scale bar, 10 μm. (B) Quantification of the number of *EGFP* RNA signals in (E). The randomly selected fields (n=6-10) of each sample were analyzed, and the RNAscope signals in approximately 400-600 cells were counted (the number of cells analyzed is indicated in parentheses on the graphs). The black horizontal lines indicate the mean of the distribution. (C) Representative, confocal z-sectioned RNAscope images of the coculture of hESCs (H9 mCh-Farneryl5) and mESCs (MS2-tagged Oct4, OM) in 2i+LIF condition. The cells were also counter-stained with Hoechst 33342 (nuclei). A confocal plane (xy) and orthogonal views (xz and yz) are shown. White arrowheads indicate the presence of mouse *Pou5f1* mRNA inside the human cells. Scale bar, 10 μm. (D) Experimental design for primed hiPSCs (GFP-positive 317D6) and naïve mESCs (tdTomato-positive J1-tdT) co-culture followed by FACS. Representative FACS analyses of mixed cells were shown either from day-5 coculture (mix with coculture) or from mix on ice prior to FACS (mix w/o coculture). GFP^+^/tdTomato^-^ and GFP^-^/tdTomato^+^ cells were sorted for the subsequent RT-qPCR analyses. (**E,F**) RT-qPCR analyses of mouse *Actb* and *Neat1* RNAs in the sorted hiPSCs from mix w/o coculture, mix with coculture, or separate culture, or of human *ACTB* and *NEAT1* RNAs in the sorted mESCs. Relative values normalized with endogenous β-actin (*ACTB*/*Actb*)

We also found mouse *Pou5f1* transcripts were detected in the acceptor hESCs upon coculture with the donor mESCs. To detect endogenously transcribed *Pou5f1* mRNAs *in situ*, we used Oct4 reporter mESCs (referred to as OM cell line), in which MS2 tag, a bacteriophage-derived stem-loop sequence, is knocked-in into the *Pou5f1* locus(13). Confocal imaging of plasma membrane reporter hESCs (H9 mCherry-Farnesyl5) showed serial z-sectioning revealed the presence of mESC-derived *Pou5f1* mRNAs inside the human cells upon coculture (**Fig. 1C**).

To further validate the inter-species mRNA transfer at the endogenously expressed genes, we used the coculture system using primed hiPSCs expressing GFP (317D6) and naïve mESCs expressing tdTomato (J1-tdT) in 2i+LIF condition for five days, followed by FACS enrichment of each cell type (referred to hereafter as “mix with coculture”) (**Fig. 1D and Fig. S2C**). To subtract the background signals potentially caused by cross contamination, we included the mixed samples of the two cell types that were grown separately, harvested, mixed on ice, and then immediately separated by the identical protocol to the coculture sample (referred to hereafter as “mix w/o coculture”) (**Fig. 1D**). Separate cultures of hiPSCs or mESCs were used as controls to determine endogenous gene expression levels before coculture. After FSC/SSC singlet cell sorting, we determined the purity of each enriched fraction by flow cytometry (**Fig. S2D**). Among at least 5,000 counted cells of each fraction, the hiPSC-enriched fraction contained no tdTomato-positive cells and vice versa. Next, we designed RT-qPCR primers and probes specific for either human or mouse β-actin mRNA (*ACTB* or *Actb*) (**Fig. S2E,F**), which have been reported to undergo transfer between mouse embryonic fibroblasts and human MCF7 breast cancer cells(3). Mouse *Actb* transcripts were more abundant in the sorted hiPSC fraction of mix with coculture condition than in the separate culture or mix without coculture condition (**Fig. 1E**). These patterns were confirmed in the other two hiPSC lines (**Fig. S2G**). The detection of mouse *Actb* transcripts in hiPSCs upon coculture is cell-contact dependent since it was significantly less detected in CM of mESC and physically separated coculture condition via transwell (**Fig. S2H**). We also found that human *ACTB* transcripts were present in the sorted mouse ESCs of mix with coculture condition (**Fig. 1F**). To test the transfer of a different type of RNAs, we evaluated a long-noncoding RNA, *NEAT1/Neat1*. Mouse *Neat1* transcripts were not detected in the hiPSCs irrespective of conditions tested, and vice versa (**Fig. 1E, F**).

The presence of inter-species mRNA transfer prompted us for comprehensively profiling by RNA-seq. We adapted the published informatics (3) to analyze the raw reads, including the length (>100 bp) and the type of fragment (paired-ends), by avoiding the mapping of mouse reads to the human genome and using data from both *in silico* simulation and sequencing experiment of the sorted cell samples. During the separation of human and mouse sequences in the obtained reads, we aligned the detected transcripts in human cells to mouse genome under the most stringent conditions to eliminate all ambiguous reads that may lead to reading misalignment. Compared to the separate culture condition, the coculture with mESCs increased the abundance of mouse-derived mRNAs found in the human cell fractions. Indeed, the percentage of mouse reads, relative to the total reads, in the hiPSC lines 317-12 and 317D6 increased from 0.05% (6,817 mouse reads) and 0.07% (8,186 mouse reads) to 0.3% (44,176 mouse reads) and 2.8% (311,859 mouse reads), respectively. On average, 9,953 transcripts of mouse genes were detected in the sorted hiPSC fractions derived from the coculture, with variable reads across genes (**Fig. S3A**). Differential gene expression analysis (fold change >2, adjusted *P* values <0.01) revealed 4,881 genes that are highly detected in the sorted human cells after mESC coculture over separate culture of hiPSCs.

We next asked if the transfer of RNAs from mESCs into hiPSCs is dependent upon the endogenous expression level in the donor cells. To explore this, we performed linear regression analysis of the mouse read enrichment in the sorted human cells from coculture (log_2_ fold change of the mouse genes of coculture over separate culture of hiPSCs) with the expression level in mESC separate culture (log_10_ mean RPKM of mouse genes). We found the overall positive correlation between the endogenous expression level and the transferred amount (Pearson correlation coefficient 0.446) (**Fig. S3B**): for example, translation factors (*Eef1b2*, *Eif2s2*), ribosomal proteins (*Rps29*, *Rpl19*, *Rps11*), and other metabolic and housekeeping genes (*Fasn*, *Scd1*, *Paics*, *Hspa8*). These indicate that the level of gene expression in donor cells reflects the intercellular RNA transfer as reported previously(1, 3).

Among the top 75 genes with the highest gene expression variance, all had higher expression in the coculture conditions (**Fig. 2A**). Gene ontology (GO) enrichment analysis revealed that a highly diverse group of genes are involved: *e.g.,* protein targeting to membrane, symbiont process, translational initiation, and RNA processing and localization (**Fig. S3C**). To analyze functional categories that are enriched in networks of the significantly abundant mouse genes, we performed over-representation analysis based on the KEGG database (**Fig. 2B**). The resultant pathways included RNA processing (RNA transport, mRNA surveillance, spliceosome, and RNA degradation), protein translation and its quality control (ribosome, protein processing in endoplasmic reticulum, ribosome biogenesis in eukaryotes, ubiquitin mediated proteolysis, and proteosome). Various signaling pathways, including pluripotency of stem cells (*Oct4, Sox2, Nanog,* and *Stat3*), metabolic reactions, cell proliferation/survival (cell cycle and survival-related growth factor signaling pathways), and cell death (apoptosis, ferroptosis, and senescence) were present. Interestingly, a series of stress-coping pathways are also enriched, including p53 signaling (35 genes including *Tp53*), DNA repair (38 genes), autophagy (71 genes), and signaling responsive to environment stresses (HIF-1 signaling, 40 genes; mTOR signaling, 73 genes; FoxO signaling, 60 genes), as well as metabolic pathways that may help stress adaptation response. In addition, *Atf4*, a transcription factor functioning as a master regulator of cellular stress responses such as endoplasmic reticulum stress and nutrient shortage, was listed as one of the most enriched transcription factor-coding mRNAs (**Table S1**). As hPSCs are exposed to 2i+LIF condition, cellular fitness might depend on such genes to thrive and adapt in an impermissible environment.

**Fig. 2.**
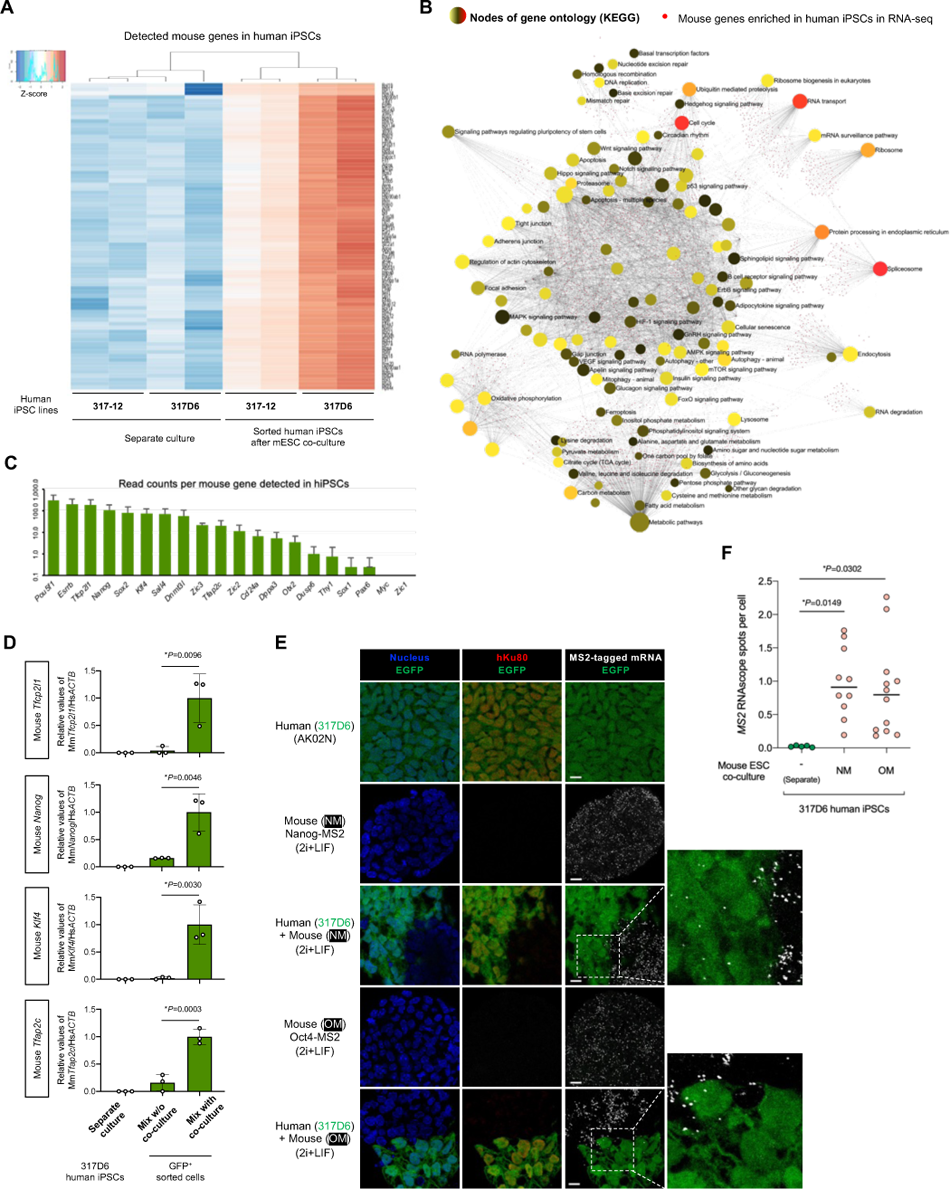
Global profiling revealed divergent mRNA transfer involving transcription factors. (A) Heatmap of differentially expressed mouse genes in hiPSCs cocultured with mESCs compared with the separate cultures of hiPSCs. The samples from two independent replicates were prepared from two hiPSC lines, 317-12 and 317D6. The top 75 genes with the highest gene expression variance across the samples were plotted. Colors represent gene expression of each sample relative to the mean. (B) Over-representation analysis (ORA) of mouse genes enriched in human the sorted hiPSCs after coculture with mESCs and the KEGG networks grouping these genes by the enrichment network tool NetworkAnalyst. The functionally grouped network was visualized based on the degree of connectivity between pathways and genes (nodes). The pathway nodes are colored according to *p*-value of ORA, while the size of the nodes corresponds to the number of genes from the analyzed gene list. The smaller nodes correspond to individual genes, colored according to their fold change. (C) Expression levels (read counts in RNA-seq) of the selected mouse transcripts related to pluripotency in hiPSCs after cocultured with mESCs. Average ± SD is shown. (D) RT-qPCR analyses of mouse *Tfcp2l1*, *Nanog*, *Klf4*, and *Tfap2c* mRNAs in the sorted hiPSCs from mix w/o coculture, mix with coculture, or separate culture. Relative values normalized with endogenous *ACTB* expression are shown as an average ± SD from three independent experiments. (E) Representative RNAscope images of MS2-tagged RNA in separate cultures of EGFP-expressing hiPSCs (317D6) in primed maintenance medium, and MS2-tagged Nanog (NM)- or MS2-tagged Oct4 (OM)-expressing mESCs in 2i+LIF, and in the coculture of hiPSCs and mESCs. The cells were also counter-stained with anti-Ku80 antibody (human nuclei) and Hoechst 33342 (all nuclei). Scale bar, 10 mm. (F) Quantification of the number of MS2 probe-positive RNA spots in (E). The randomly selected fields (n=6-12) of each sample were analyzed for the RNAscope spots in over 500 cells. The black horizontal lines indicate the mean of the distribution.

Recently Hu et al reported that the RNA sensing pathway called retinoic acid-inducible gene I (RIG-I) pathway plays an important role in the interspecies PSC competition which determines the winner cell states at the primed pluripotency(14). They utilized the coculture of human and mouse primed PSCs and found the existence of cell contact-dependent human-to-mouse RNA transfer, which activates the RIG-I pathway in the surviving mouse cells. We asked if the RIG-1 pathway is upregulated in hPSCs upon coculture with naïve mESCs by analyzing the RNA-seq of human cells sorted from the coculture. The surviving hiPSCs did not upregulate but rather downregulated the RNA receptors *DDX58* (RIG-I) and *IFIH1* (MDA5) without evident activation of the downstream genes such as *IRF7*, *IFITM3*, *IFIT1*, *ISG15*, and *ISG20* (**Fig. S3E**). Among the mouse mRNA detected in the coculture conditions, we found 491 genes related to transcriptional regulator activities (**Table S1**). The pathway enrichment analysis revealed that signaling pathways regulating pluripotency of stem cells were among the over-represented pathways (ranked sixth) (**Fig. S3D**). These factors regulating pluripotency are clustered into the core network (*Nanog, Pou5f1, Esrrb*, and *Sox2*), and the downstream signaling of SMAD family protein (*Smad2, Smad3*, and *Smad5*) and Wnt signaling (*Ctnnb1* and *Tcf3*) (**Fig. S3F**). This is consistent with the finding that the detected set of mouse-derived transcription factor mRNAs includes *Pou5f1, Esrrb, Tfcp2l1, Nanog, Sox2*, and *Klf4* as transcripts with abundant counts in the RNA-seq (**Fig. 2C**). We validated the RNA-seq results by RT-qPCR and RNAscope analyses. Compared with the mix without coculture condition, the direct contact dependent coculture with mESCs (mix with coculture) led to the pronounced transfer of mouse-derived *Tfcp2l1, Nanog, Klf4*, and *Tfap2c* mRNAs into human iPSCs as determined by RT-qPCR of FACS sorted hiPSCs from the coculture (**Fig. 2D**, **Fig. S2G**, and **Fig. S4A, B**). To detect the transfer of endogenous transcription factor-coding mRNAs *in situ*, we cocultured hiPSCs (EGFP-positive, 317D6 line) with Nanog or Oct4 reporter mESCs (referred to as NM or OM cell line, respectively), in which MS2 tag, a bacteriophage-derived stem-loop sequence, is knocked-in into the respective gene loci(13, 15) (**Fig. S4C**). Using the MS2-specific RNAscope probe, the transfer of mouse *Nanog* and *Oct4* mRNAs were detected within hiPSCs in the mouse ESC coculture, which displayed significantly more abundant RNAscope signals than those under separate culture condition, and no signals were detectable in the negative control probe staining (**Fig. 1C**, **Fig. 2E, F** and **Fig. S4D, E**). These results indicate that transcription factor-coding mRNAs are transferred from mouse ESCs into human PSCs during the coculture.

Given the inter-species transfer of various mRNAs in PSCs in a cell-contact dependent manner, we examined the physical contact sites in the coculture of primed hESCs and naïve mESCs. Consistent with previous observations of cell contact-mediated mRNA exchange(1, 3, 12, 16), we observed thin protrusions that appeared to connect hPSCs and mESCs in 2i+LIF condition (**Fig. 3A**). The structures were positive for both membrane-specific dye and F-actin, most of which were extended from hESCs as revealed by the expression of membrane reporter mCherry-Farnesyl5. In RNAscope assay we found the presence of *EGFP* mRNA in the protrusion of hESCs when cocultured with EGFP reporter mESCs (**Fig. 3B**). We also found that hESCs extend their membranes toward the mESCs near the bottom of coculture over the distance corresponding to 2-3 cells (**Fig. 1C** and **Fig. 3B**), and that these extensions contain mESC-derived mRNAs as assessed by visualization of MS2-tagged *Pou5f1* mouse RNAs (**Fig. 3C**).

**Fig. 3.**
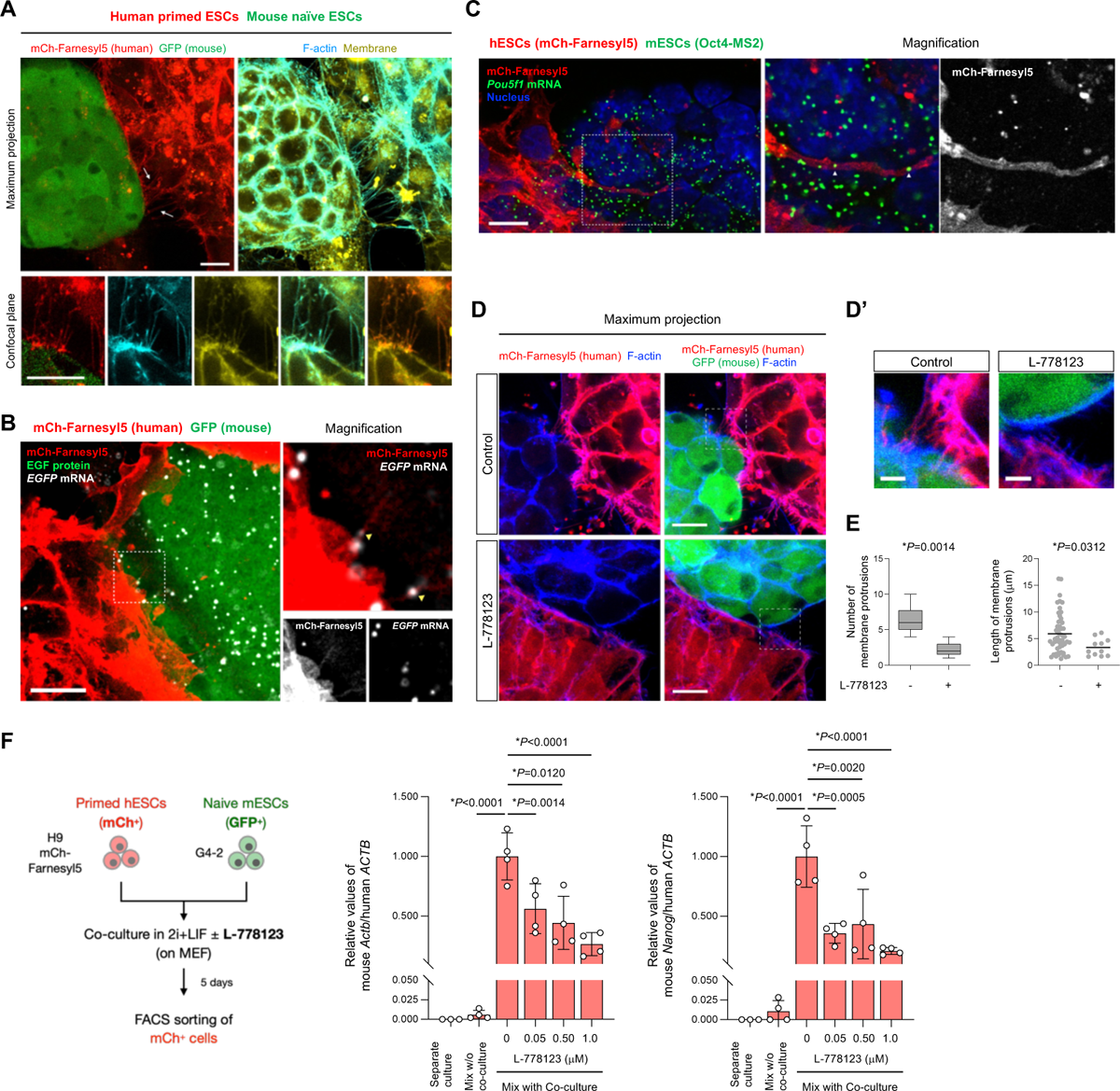
mRNA-containing membrane protrusions from mouse to human cells are required for mRNA transfer. (A) Co-culture of H9 mCherry-Farnesyl5 (red) hESCs and G4-2 GFP (green)-expressing mESCs stained with F-actin (cyan) and membrane (yellow) specific dyes. Top images are maximum projection of z-stacked images. Bottom images are from a confocal plane of a magnified region containing membrane protrusions. Arrows indicate membrane protrusions connecting hESCs and mESCs. (scale bar, 10 mm) (B) Representative confocal images of *EGFP* RNA (white) in the coculture of H9 mCherry-Farnesyl5 (red) hESCs and G4-2 GFP (green)-expressing mESCs visualized by RNAscope (scale bar, 10 mm). The area indicated by the dashed box is also magnified in the right panels. (C) Representative confocal images of MS2-tagged *Pou5f1* RNA (green) in the coculture of H9 mCherry-Farnesyl5 (red) hESCs and Oct4-MS2 mESCs visualized by RNAscope (scale bar, 10 mm). The area indicated by the dashed box is also magnified in the right panels. (**D-D’**) Maximum projection images of the co-culture of H9 mCherry-Farnesyl5 (red) hESCs and G4-2 GFP (green)-expressing mESCs treated either with control or with L-778123 (0.5 mM). The cells were counterstained with F-actin-specific dye (blue) (D, scale bar, 10 mm). The area indicated by the dashed box is also magnified in D’ as a confocal plane (scale bar, 5 mm). (E) Quantification of the number (left) and the length (right) of membrane connections observed between hESCs and mESCs in the presence or absence of L-778123. (F) Experimental design for primed H9 hESCs (mCherry-positive) and naïve G4-2 mESCs (GFP-positive) co-culture in the presence of L-778123 followed by FACS sorting of mCherry-positive fraction (left, also see **Fig. S5B**). RT-qPCR analyses of mouse *Actb* and *Nanog* RNAs in the sorted hESCs from mix with co-culture conditions treated with the indicated concentrations of L-778123. The separate culture as well as mix w/o co-culture was set as control. Relative values normalized with endogenous human *ACTB* expression are shown.

Intercellular tunneling nanotube-like structures have been implicated in mediating the transfer of various cellular materials including RNAs and organelles(17–21). The membrane remodeling associated with F-actin is regulated by GTPase family proteins, in which the prenyl group (farnesyl or geranylgeranyl) modification are essential for their membrane association and biological activities. To target the membrane protrusions observed in the coculture of hESCs and mESCs, we treated cells with L-778123, a type 1 farnesyltransferase and geranylgeranyltransferase inhibitor(22). Upon L-778123 treatment the mCherry-Farnesyl5 reporter, which contains the farnesylation site, lost its membrane localization and diffused into cytoplasm in hESCs (**Fig. S5A**). L-778123 significantly reduced the formation of membrane protrusions and their length observed in the contact site between hESCs and mESCs (**Fig. 3D,E**). To investigate the effect of L-778123 on the intercellular mRNA transfer, we isolated the human cells based on mCherry expression from L-778123-treated, mESC coculture followed by RT-qPCR analysis of mouse *Actb* and *Nanog* genes (**Fig. S5B**). L-778123 treatment upon coculture decreased the detected levels of mouse *Actb* and *Nanog* transcripts in the sorted hESCs (**Fig. 3F**). In this situation, the expression of the two genes as well as pluripotency markers in mESCs remain unchanged (**Fig. S5C**). Collectively, these results indicate that the mESC-derived mRNAs can be delivered into the neighboring hESCs at least in a part via intercellular membrane protrusions.

To explore the possibility that the donor cell-derived mRNAs are translated in the acceptor cells, we measured the reporter fluorescence protein signals of acceptor human cells in the coculture of GFP-expressing primed hiPSCs and tdTomato-expressing naïve mESCs described in **Fig. 1D**. While we did not capture significant tdTomato signals inside human cells under confocal microscope, which might be due to the detection limit in our microscope setup, in flow cytometry analysis we found the increased shift of mean tdTomato intensity in mix with co-culture compared to separate culture and mix without co-culture conditions (**Fig. S6A**). In the membrane connections between hiPSCs and mESCs, we did not observe the EGFP signals in mCherry-Farnesyl5-positive protrusions (**Fig. 3A**), indicating less possibility of direct EGFP protein transfer. We also assessed if endogenous genes are detected in protein levels in human cells upon coculture with mESCs. We used mouse-specific antibodies against Oct4 and Nanog. Using separate culture of human iPSCs and mouse ESCs, we validated the specificity of these antibodies (**Fig. S6B**), which never cross-reacted with human orthologues. Upon coculture with mESCs, the human cells in close proximity to mESCs were dim-positive for mouse Oct4 and Nanog as revealed by the mouse-specific antibody immunostaining (**Fig. S6C,D**). These results indicate the presence of mouse transcription factor proteins inside human cells upon coculture with mESCs.

Cellular reprogramming can be achieved by the combinatorial introduction of mRNA subsets (23) with broad transcription control ability(24, 25). Given that mRNA transfer involves transcription factors, we hypothesized that coculture might influence cellular phenotype. Relative to mESC in a naïve state, conventional hPSCs, either ESCs or iPSCs, represent a more developmentally advanced, morphologically distinct, primed pluripotent state. We found that hPSCs after five days of naïve mESCs coculture in 2i+LIF condition underwent naïve-like morphological change (**Fig. 4A, B** and **Fig. S1F**). This naïve-like morphology became more prominent after the sorting human cells from the coculture and the subsequent culture in PXGL, a medium for culturing naïve hPSCs(26), confirmed by the presence of hESCs (H9) with dome-shaped morphology (**Fig. 4A**). Several other hiPSC lines (317-12, 317D6, and TkDA) also reproduced naïve-like morphological conversion after the coculture with mESCs (**Fig. S7A**). RT-qPCR analyses of the hiPSCs after human cell isolation revealed the upregulation of naïve pluripotency markers (*DPPA3, TFCP2L1, DNMT3L, KLF4,* and *KLF17*) and downregulation of primed marker (*DUSP6*) compared with the separately cultured cells in primed states (**Fig. S7B**). We also found with flow cytometry that five to six days after sorting following the coculture with mESCs, a proportion of human H9 cells (3–4%) were positive for SUSD2, a specific surface marker of human naïve PSCs(27, 28), as was seen in positive control chemically reset (cR) cells(26, 27) established from the same cell line (**Fig. 4C**). In contrast, most H9 cells cultured in 2i+LIF without m ESCs were mostly SUSD2-negative, similar to conventional, primed cells. We detected other cell surface markers of naïve hPSCs such as CD130 and CD77 in hiPSCs cocultured with mESCs (**Fig. S7C**). In contrast, the expression of CD90 and HLABC reported as primed hPSCs markers (29) decreased.

**Fig. 4.**
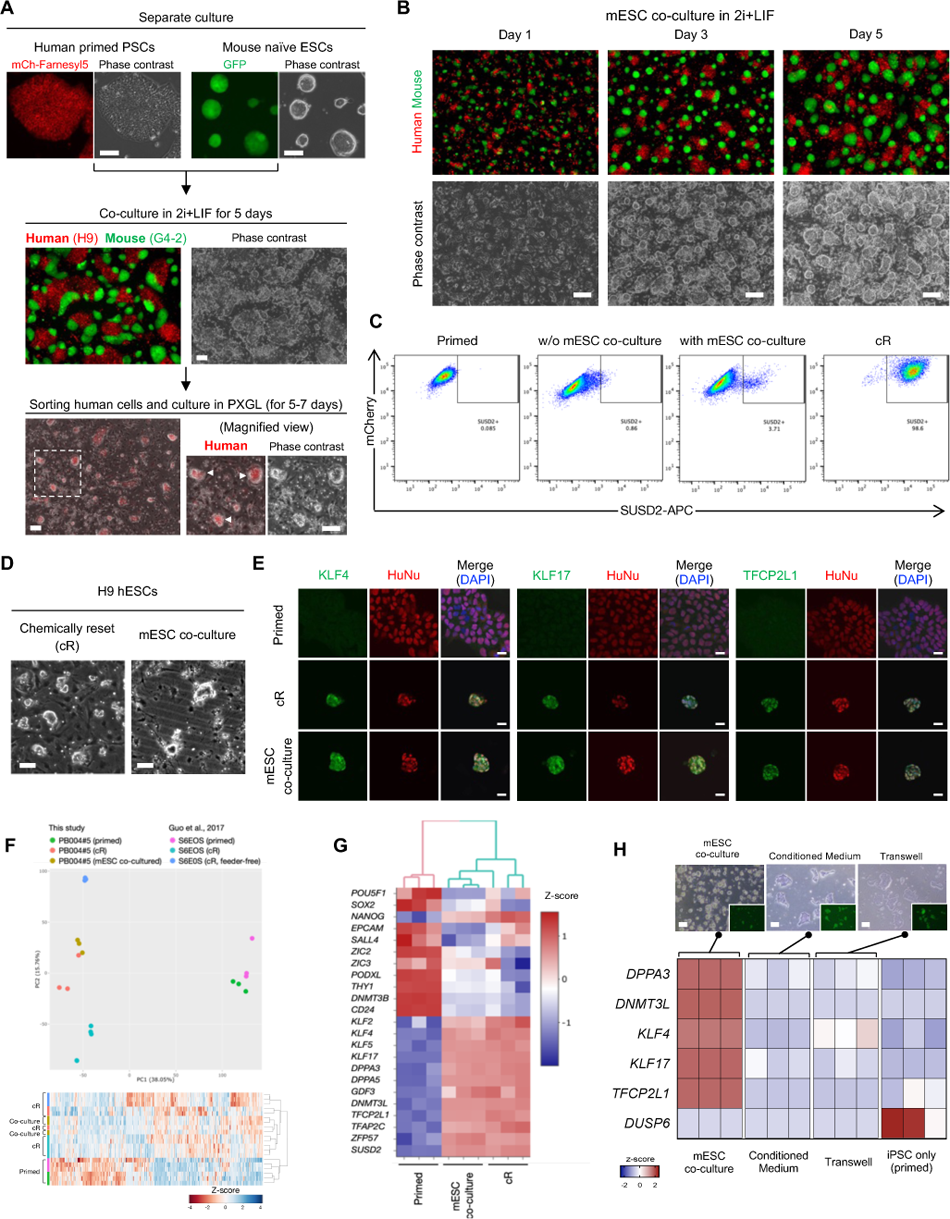
Inter-species coculture with mouse ESCs enables naïve-like conversion of human PSCs. (A) Protocols for coculture of primed hPSCs (H9 human ESCs expressing a membrane reporter mCherry-Farnesyl5) and naïve mESCs (G4-2 mouse ESCs expressing GFP). Both cell types (50% each) were cocultured in 2i+LIF condition for 5 days followed by sorting human cells. The sorted human cells were further cultured in PXGL condition, representing the morphological conversion into dome-shaped colonies (white arrowheads). Scale bar, 100 mm. (B) Representative images captured at the indicated time points during the coculture of primed hPSCs (H9, red) and naïve mESCs (G4-2, green). Scale bar, 100 mm. (C) Flow cytometry analysis using naïve-specific SUSD2 antibody in H9 hESCs (expressing mCherry) that were separately cultured in a primed state (primed) or 2i+LIF followed by culture in PXGL (w/o mESC coculture), or cocultured with mouse ESCs in 2i+LIF followed by human cell sorting and culture in PXGL (with mESC coculture). The chemically reset cells (cR) derived from the same H9 cells were also included. (D) Phase-contrast images in H9 hESCs that were converted into cR cells (left) or expanded after sorting SUSD2^+^ cells from the cocultured H9 cells with mESCs (right). Scale bar, 100 mm. (E) Immunostaining images of human naïve pluripotency markers KLF4, KLF17, and TFCP2L1 together with human-specific nuclear antigen (HuNu) in parental primed hiPSCs (upper), cR cells (middle), and SUSD2^+^-sorted human cells cocultured with mESCs (bottom). Scale bar, 20 mm. (F) Principal-component analysis (PCA) was performed on RNA-seq measurement obtained from hiPSCs, including primed cells (parental), cR cells, and converted cells via mESC coculture and subsequent expansion. The data were clustered with previous, independently generated RNA-seq of hESCs in primed and cR states. (G) Unbiased hierarchical clustering of samples represented in (**F**). The expression levels of selected naïve and primed pluripotency marker genes are shown. (H) Expression heatmap of selected naïve and primed pluripotency markers in hiPSCs, including direct coculture with mESC, conditioned medium from mESCs, transwell coculture, and primed cells. The representative images of each condition are shown (top, scale bar, 100 mm).

We next asked how native gene expression of hPSCs is impacted by coculture with mESCs after human cell sorting and subsequent culture in PXGL. We performed RNA-seq at different timepoints during mESC coculture-dependent phenotypic conversion. Pattern clustering revealed seven distinct clusters of genes, which include down-regulation of genes related to cell adhesion molecules and neuroactive ligand-receptor interaction toward the converted state as well as up-regulation of genes related to signaling pathways regulating pluripotency of stem cells (**Fig. S7D**). We observed the transient up-regulation of genes during the coculture with mESCs that function in calcium signaling pathway and focal adhesion. Since cell-cell and cell-matrix adhesion interactions are dynamically rearranged between naïve and primed pluripotency and intracellular calcium contributes to these regulations(28, 30, 31), the observed transitioning transcriptome may influence the human cells adapting to the culture environment for naïve PSCs. While a portion of human cells maintained the expression of Oct4 and Nanog upon coculture with mESCs in 2i+LIF (**Fig. S1A-D**), we found a subset of naïve pluripotency signature genes, such as *TBX3*, *IL6ST*, *TFAP2C*, and *DMNT3L*, being expressed in the coculture period at the intermediate levels between primed and fully converted states (**Fig. S6E**). Expression levels of some genes of pluripotency signaling including *KLF2* and *LEFTY2* are comparable with those in converted cells and cR cells (**Fig. S7D,E**). These data indicate that hPSCs adopt to the state close to the naïve pluripotency after sorting from coculture with mESCs.

To further characterize the converted cell phenotype induced by the co-culture, the SUSD2-positive sorted cultures of human cells were maintained for several passages (**Fig. 4D** and **Fig. S1F**). We carried out immunofluorescence analysis in the propagated putative naïve hiPSCs. Consistent with cR cells, co-culture-derived hiPSCs expressed naïve markers KLF4, KLF17, and TFCP2L1 with human specific nuclear antigen, whereas parental, separately cultured primed hiPSCs did not (**Fig. 4E**). Unbiased clustering and dimension reduced plot analysis of transcriptional profiles of RNA-seq data demonstrated that cR cells and the propagated cells via mESC co-culture share nearly indistinguishable gene expression patterns, while both profiles were significantly different from those of the primed hiPSCs (**Fig. 4F**). The transcriptome of mESC-co-cultured cells showed high correlation with that of cR cells both in-house and in previously published, extended passage cell datasets(26) (**Fig. S8A**). The naïve markers including key transcription factors (*KLF2, KLF4, KLF5, KLF17, TFCP2L1*, and *TFAP2C*), epigenetic regulators (*DPPA3, DPPA5*, and *DNMT3L*), and transposable element regulator (*ZFP57*) were highly expressed in cR and mESC-co-cultured cells (**Fig. 4G**). Conversely, these cells downregulated primed pluripotency markers including *THY1, DNMT3B*, and *CD24* as well as lineage commitment genes like *SALL4, ZIC2/ZIC3*, and *PODXL* compared to the primed hiPSCs. Compared with the primed hiPSCs, the cR and mESC-co-cultured cells displayed 5,195 and 5,467 differentially expressed genes, respectively, with a high level of shared genes between the two conditions (**Fig. S8B,C**). Conditioned medium from mESCs and transwell coculture assays could not induce dome-shaped hiPSC phenotype and resulted in negligible induction of naïve pluripotency markers (**Fig. 4H**). The observed down-regulation of *DUSP6*, a primed marker gene that is induced by FGF-stimulated Erk signaling, can be attributed to an inhibition of Erk by PD0325901, a component of 2i+LIF and PGXL media.

We also studied the loss of imprinting of mESC co-culture-induced converted hPSCs, a hallmark of naïve, but not primed, PSCs, by studying biallelic expression genes in RNA-seq data (see experimental procedures). By calling single-nucleotide variants in certain exons, we listed imprinting genes which were expressed biallelically in either our sample while referring to the previous, large-scale analysis of imprinting in human PSCs(32, 33). In line with these studies our cR naïve hESCs as well as mESC co-culture-induced converted cells represented loss of imprinting in *ZDBF2*, *IGF2*, *DLK1*, and *ZNF597* while primed cells did not (**Fig. S8D**), indicating the similar pattern of loss of imprinting in hPSCs along the mESC co-culture-induced conversion as conventional chemical reset naïve state. Collectively, these results indicate that direct cell contact with mESCs induced the phenotypic conversion of primed hPSCs into naïve state.

To extend our observation to other interspecies conditions, we used the rat naïve iPSC line (riPSC), which has been generated in our previous reports(34, 35). Similarly to the co-culture with naïve mESCs, the riPSCs constitutively expressing EGFP were co-cultured with primed H9 hESCs expressing mCherry-Farnesyl5 under 2i+LIF condition for 5 days (**Fig. S9A**), followed by FACS sorting of mCherry^+^ EGFP^-^ and mCherry^-^ EGFP^+^ cells (**Fig. S9B**). RT-qPCR of the sorted mCherry^+^ EGFP^-^ human cell fractions using rat-specific primers and probes revealed the significant amount of rat-derived transcripts, such as *Actb*, *Klf4*, *Nanog*, *Tfap2c*, and *Tfcp2l1*, detected in the mix with coculture condition compared with the separate culture or mix without coculture condition (**Fig. S9C-E**). The human *ACTB* was present in riPSCs upon mix with co-culture, suggesting the intercellular, bidirectional RNA exchange between these cells (**Fig. S9E**). After the human cell sorting and subsequent culture in PXGL, the SUSD2^+^ human cells emerged from both conditions of co-culture with mESCs and riPSCs, as revealed by flow cytometry (**Fig. S9F**). We further expanded the SUSD2^+^ cells in PXGL that displayed dome-shaped morphology (**Fig. S9G**). RT-qPCR of the expanded cells showed the upregulation of naïve marker genes and the downregulation of a primed marker gene *DUSP2* (**Fig. S9H**). These results suggest that the interspecies mRNA transfer and subsequent naïve-like conversion of hPSCs also occur in the other context of co-culture with rat naïve pluripotent stem cells.

Next we examined if the human cells that receives RNAs from the donor mESCs transition to a naïve-like state. In order to track the distribution of donor-derived RNAs during co-culture, we first transfected Cy5-labelled GFP mRNA as a tracer mRNA into naïve mESCs and then co-cultured them with primed hESCs (H9 mCh-Farnesyl5) in 2i+LIF for 5 days (**Fig. 5A**). Flow cytometry of the day 5 co-culture cells revealed that mCherry-negative mESCs contained both Cy5 and GFP signals (**Fig. 5B**). A portion of hESCs also contained Cy5 signals upon co-culture with Cy5-mRNA transfected mESCs, indicating the intercellular transfer of the tracer mRNA from mESCs into hESCs (**Fig. 5B**). The sorted Cy5-positive human cells (mCherry-positive) more expanded in the PXGL medium with the shape of domed-like colonies than Cy5-negative cells (**Fig. 5C**). Consistent with this, the Cy5-positive human cells expressed higher levels of a proliferation marker gene *MKI67* than Cy5-negative cells (**Fig. 5D**). To assess the efficiency of conversion into naïve-like state, we performed flow cytometry of the naïve PSC surface marker SUSD2 in the expanded human cells. Compared to the Cy5-negative cells, the Cy5-positive human cells consisted of higher proportion of SUSD2 expressing cells 5 days after the human cell sorting (day 10 after the start of co-culture) (**Fig. 5E**). RT-qPCR analysis verified the more naïve-like induction in Cy5-positive human cell fraction which expressed higher levels of naïve markers *NANOG*, *KLF4*, *TFCP2L1*, and *TFAP2C* as well as lower expression of the primed marker *DUSP6* (**Fig. 5F**). The treatment with L-778123 during the co-culture inhibited the induction of naïve marker genes *NANOG*, *KLF4*, and *TFAP2C* following isolation of human cells and subsequent culture in PXGL (**Fig. 5G**). These results support the notion that the hPSCs receiving mRNAs from mESCs are prone to convert into naïve-like state.

**Fig. 5.**
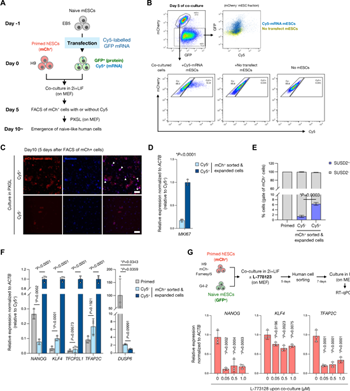
mRNA transfer-prone human PSCs efficiently convert into naïve-like state. (A) Experimental design for tracing a reporter mRNA transfected in mESCs during the coculture in 2i+LIF condition. *In vitro* transcribed, Cy5-labelled GFP mRNA was transfected in mESCs 1 day before the start of coculture with primed hESCs (mCherry-positive). Day 5 post coculture, mCherry-positive human cells were sorted into either Cy5-positive or -negative fraction. (B) Representative FACS analyses of the coculture at day 5. In mCherry-negative fraction, both Cy5 and GFP signals were higher in the coculture with Cy5-GFP mRNA transfected mESCs compared to that with non-transfected mESCs (top panels). To gate the Cy5-positive human cells, coculture condition with non-transfected mESCs as well as one without mESCs (hESC only) was set as controls (bottom panels). (C) Expansion of Cy5-positive and -negative hESCs (mCherry-positive) in PXGL medium after isolation from coculture with Cy5-GFP mRNA transfected mESCs. Scale bar, 200 mm. (D) RT-qPCR analysis of a proliferation marker *MKI67* of the cells in (**C**). Average ± SD is shown from three independent experiments. (E) Flow cytometry analysis of a naïve PSC surface marker SUSD2 of the cells in (C). Primed hESCs were set as a control. (F) RT-qPCR analyses of naïve or primed marker genes expressed in the cells in (C). Primed hESCs were set as a control. Relative values normalized with endogenous *ACTB* expression are shown as an average ± SD from three independent experiments. (G) Experimental design for primed H9 hESCs (mCherry-positive) and naïve G4-2 mESCs (GFP-positive) co-culture in the presence of L-778123 followed by FACS sorting of mCherry-positive fraction and expansion in PXGL medium (top). RT-qPCR analyses of *NANOG*, *KLF4*, and *TFAP2C* expression in the expanded hESCs from mESC co-culture.

To examine the causal relationship of mESC-derived transferred mRNA on hPSC conversion into naïve-like state, we established a loss-of-function system in which mouse-derived transcripts in hiPSCs are specifically downregulated via shRNAs targeting mouse transcription factors (**Fig. 6A**). Among the list of mouse-derived transcription factors identified in the RNA-seq (**Fig. 2C**), we focused on *Tfcp2l1, KLF4,* and *Tfap2c* genes, since they are essential for the maintenance of human naïve pluripotency while self-renewal in the primed state is not affected when these genes are inactivated(36, 37). Based on the alignment of human and mouse transcripts for each gene, we designed shRNA sequences to target mouse, not human, genes (**Fig. S10A**), followed by packaging them into lentivirus. To validate the specificity and efficacy of the designed shRNAs, we transduced these lentivirus-packaged shRNAs into independently propagated mESCs and hiPSCs in a separate culture and selected shRNAs that specifically down-regulated mouse *Tfcp2l1/Klf4/Tfap2c* genes (**Fig. 6B**; the shRNAs used are shKlf4 #1, shKlf4#3, shTfcp2l1 #1, shTfcp2l1 #2, shTfcp2l1 #3, and shTfap2c #3). We next generated hiPSCs (Venus-positive) stably expressing the selected shRNAs and confirmed that shRNA expression affected neither Oct4 expression in primed states nor naïve conversion efficiency via chemical resetting (**Fig. S10B, C**). After the coculture of shRNA-expressing hiPSCs with mESCs in 2i+LIF condition, we passaged the cells and continued the culture in the presence of puromycin to select human cells (**Fig. 6A**). As a control, luciferase-targeting shRNA expression resulted in the emergence of domed shape colonies after the puromycin selection while knockdown of mouse-derived *Klf4, Tfcp2l1*, and *Tfap2c* in hiPSCs induced no or few colonies (**Fig. 6D**). Immunostaining a naïve specific marker KLF17 in the remnant colonies after puromycin selection demonstrated that, unlike control shRNA, all tested shRNAs targeting mouse *Tfcp2l1, Klf4* and *Tfap2c* did not produce KLF17-positive cells (**Fig. 6E**). Together, inactivation of mouse-derived *Tfcp2l1, Klf4,* and *Tfap2c* transcription factor-coding mRNAs in hPSCs abolishes the naïve reprogramming potential induced by mouse naïve PSC coculture.

**Fig. 6.**
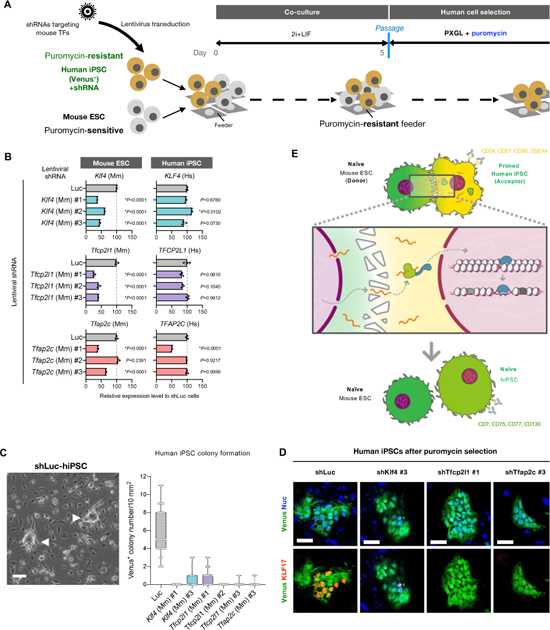
Pioneer transcription factor mRNA transfer is indispensable for naïve conversion of hiPSCs. **(A)** Protocols for coculture of primed hiPSCs expressing Venus and shRNA specifically targeting each mouse transcription factor (puromycin-resistant) with naïve mESCs (puromycin-sensitive) followed by the induction of naïve-like conversion. After the coculture of both cell types (50% each) in 2i+LIF condition for 5 days, they were passed onto the puromycin-resistant feeder cells in the presence of puromycin in PXGL medium to eliminate mESCs. After the puromycin selection, the residual human cells were subsequently analyzed. **(B)** RT-qPCR analysis of endogenous *Klf4, Tfcp2l1, Tfap2c, Nanog*, and *Pou5f1* gene expression levels in mESCs in which lentiviral shRNAs targeting mouse *Klf4, Tfcp2l1, Tfap2c, Nanog*, or *Pou5f1* designed in **fig. S9A** were expressed. **(C)** RT-qPCR analysis of endogenous *KLF4, TFCP2L1, TFAP2C, NANOG*, and *POU5F1* gene expression levels in hiPSCs in which lentiviral shRNAs targeting mouse Klf4, Tfcp2l1, Tfap2c, Nanog, or Pou5f1 designed in **fig. S9A** were expressed. **(D)** The emerging colonies of dome-shaped, human cells expressing control luciferase shRNA (shLuc) emerging after puromycin selection is shown (white arrowheads in the left image; scale bar, 100 mm). The number of Venus-positive, dome-shaped colonies emerging after the puromycin selection in each shRNA expression condition was shown in the box and whiskers plot (right; bars represent min to max of the counts). **(E)** Immunostaining images of human naïve pluripotency marker KLF17 in Venus-positive hiPSCs expressing shRNAs targeting luciferase (shLuc) or mouse *Klf4, Tfcp2l1*, and *Tfap2c* after the coculture with mESCs followed by puromycin selection. Scale bar, 50 mm. **(F)** Schematic representation of mESC co-culture-driven reprogramming of primed hPSCs into naïve state via mESC-derived transfer. During the coculture, mRNAs that plays pleiotropic roles including stress responses are transferred between the two cells in a manner dependent on the direct cell contact.

RNA transfer via cell contact dependent (3, 38) or independent (39, 40) mechanism is gaining attention. Our experimental human and mouse pluripotent stem cell culture system uncovered the presence of extraordinary mRNA transfer that can influence neighboring cellular phenotype. Transfer specific global profiling indicates that the mobile mRNA transfer (i) is dependent on direct cell-to-cell contact, (ii) contains divergent stress-coping genes, including ER stress protection genes, such as *ATF4* and apoptosis inducing-genes such as *p53*, (iii) elicits both cellular “cooperative” and “competitive” events in the receivers, the part of which is found in interspecies PSC coculture (4), and (iv) involves a readily reprogrammable amount of transcription factor genes, which results in alternations of the acceptor cell fate. These findings indicate that mRNA transfer would permit the development of genetically and chemically neutral reprogramming approaches without nuclear transfer (41–43), cell fusion (44–46), transcription factor gene transduction (24, 47, 48), or small molecules (49, 50). The biological significance of the mRNA transfer mechanism sets the stage for further investigation in broadening the view of intra- and inter-species cell communication.

## ACKNOWLEDGMENTS

The authors would like to express their sincere gratitude to Dr Vivian Hwa for critical reading of the manuscript, Dr. Yuya Ogawa and Dr. Knut Woltjen for kindly providing mouse ESCs and GFP-expressing human iPSC lines 317-12 and 317-D6, Dr. Laurent David for providing us mouse-specific and human-specific NANOG primers, and technical assistance; Dr.Yueh-Chiang Hu for kindly providing EGFP mESC lines. We thank M. Kofron for very supportive and kind instructions on using confocal microscope and Imaris8. We also thank Drs. James M. Wells, Aaron M. Zorn, Leah Kottyan, Rafi Kopan and Stephen Waggoner for their helpful comments on this project, and all CuSTOM affiliated lab members at CCHMC. This work was supported by Cincinnati Children’s Research Foundation grants to Drs. Takebe and Weirauch, AMED grants 23bm1223006h0001, JP18fk0210037h0001, JP18bm0704025h0001, JP21gm1210012h0002, JP21bm0404045h0003, and JP21fk0210060h0003, JST Moonshot JPMJMS2022-10 and JPMJMS2033-12, and JSPS KAKENHI Grant JP18H02800, 19K22416. This project was partly supported by PHS Grant P30 DK078392 (Integrative Morphology Core) of the Digestive Disease Research Core Center in Cincinnati and by the Falk Transformational Awards Program, NIH DP2 DK128799-01, R01DK135478 and CREST (20gm1210012h0001) grant from Japan Agency for Medical Research and Development (AMED) to TT.

## Methods

### Culture of PSC lines

Human iPSC lines 1383D6 (RIKENBRC #HPS1006, obtained from RIKEN cell bank of RIKEN BioResource Research Center (BRC)) (51), 317-12 and 317D6 (kindly provided by K Woltjen, Kyoto University)(52), TkDA3-4 (obtained from The Institute of Medical Science, The University of Tokyo) (53), and PB004(54), which were derived from peripheral blood mononuclear cells or skin fibroblasts of healthy donors, which were deidentified prior to use in this study. The human primed iPSC lines as well as human primed ES cell line H9 (#WA09, obtained from WiCell Research Institute) were maintained without feeder cells either in mTeSR1 (STEMCELL Technologies) on growth factor reduced Matrigel (Corning) coated dishes or plates, or in StemFit AK02N (Ajinomoto) on iMatrix-511 (Nippi) coated dishes or plates. The primed PSCs were passaged using Accutase (Innovative Cell Technologies) every 7 days.

Mouse ESC lines EB5 and G4-2 (RIKEN CELL BANK), and J1 (ATCC) were maintained without feeder cells on dishes coated with gelatin (Sigma-Aldrich), or with poly-L-ornithine (FUJIFILM Wako Pure Chemical) and laminin (FUJIFILM Wako Pure Chemical) in N2B27 medium containing 1 μM PD0325901 (Selleck), 3 μM CHIR99021 (FUJIFILM Wako Pure Chemical), and 1,000 units/ml mouse leukemia inhibitory factor (LIF; Millipore). The N2B27 medium consisted of 1:1 mixture of DMEM/F12 (Sigma-Aldrich) and Neurobasal (Gibco), 0.5× N2-Supplement (Gibco), 0.5× serum free B27-Supplement (Gibco), 1× Glutamax Supplement (Gibo), 1× monothioglycerol (FUJIFILM Wako Pure Chemical), 0.05% bovine albumin fraction V (Gibco) and 1× non-essential amino acid (Gibco). The G4-2 mouse ESC line was derived from EB3 and carried the CAG promoter-driven EGFP(55). The *Nanog* and *Oct4* MS2 reporter mouse ESC lines, NM and OM, were described previously(13, 15). Cell cultures were regularly checked for mycoplasma contamination.

Rat iPSC line WI-T1-3 generated from Wistar rat embryonic fibroblast by introducing three mouse reprograming factors (Oct3/4, Klf4, Sox2) in a lentiviral vector as a doxycycline-inducible expression, which also contains EGFP driven by Ubc promoter(34, 35). The riPSCs were maintained on MEF feeders in N2B27 medium containing 1 μM PD0325901, 3 μM CHIR99021, and 1,000 units/ml mouse leukemia inhibitory factor. Upon passage Matrigel was added at the dilution of 1:100 into the medium.

Chemical conversion to human naïve PSCs was performed as previously described(26, 27). Primed human PSCs (1×10^4^ cells/cm^2^) were seeded on mouse embryonic fibroblast (MEF) feeder cells in AK02N with 10 μM Y-27632 (nacalai tesque) and cultured. The following culture was performed under 5% O_2_ condition. After overnight incubation, the medium was replaced with Prime ES Cell Medium (ReproCELL) supplemented with 5 ng/ml basic FGF (nacalai tesque) followed by culture for 24 hr under 5% O_2_ condition. The medium was then switched to cR-1 (N2B27 + 1 μM PD0325901, 20 ng/ml human LIF (Peprotech), and 0.75–1.0 mM valproic acid sodium salt (VPA; Sigma-Aldrich)). After 3 days, the medium was switched to PXGL (N2B27 + 1 μM PD0325901, 20 ng/ml human LIF, 2 μM XAV939 (FUJIFILM Wako Pure Chemical), and 2 μM Gö6983 (Selleck)). After further culture for around 7 days, cells were passaged with Accutase on MEF feeder cells in PXGL. To enrich naïve PSCs, cells were detached with Accutase followed by live cell staining with APC-conjugated SUSD2 antibody (BioLegend) and Ghost Dye Violet 450 (TONBO) on ice for 30 min. The SUSD2^+^ cells were sorted either by fluorescent-activated cell sorting (FACS) on FACS Aria3 (BD Biosciences), or by magnetic-activated cell sorting (MACS) with anti-APC MicroBeads (Miltenyi Biotec) on MACS MultiStand separator (Miltenyi Biotec) according to the manufacturer’s instruction. The emerging dome-shaped colonies were passaged every 5-7 days and maintained in PXGL.

### Coculture of human and mouse/rat PSCs

Coculture was performed under 5% O_2_ condition. Primed human PSCs and naïve mouse ESCs/rat iPSCs were dissociated into single cells with Accutase, and total 2–4×10^5^ cells (1–2×10^5^ cells each for human or mouse cells) were seeded on MEF feeder cell-coated 12-well plate in N2B27 medium containing 1 μM PD0325901, 3 μM CHIR99021, 20 ng/ml human LIF, and 10 μM Y-27632. On the next day (day 1), the medium was replaced with N2B27 medium containing 1 μM PD0325901, 3 μM CHIR99021, and 20 ng/ml human LIF. The medium was replaced every day till day 5. On day 5, mixed cells were treated with Accutase into single cells followed by sorting human cells either by FACS or by MACS. For MACS, mouse cell depletion kit (Miltenyi Biotec) was used to enrich human cells. Sorted human cells (1×10^5^ cells/well) were seeded on MEF feeder cell-coated 12-well plate in PXGL (27) containing 10 μM Y-27632 and Geltrex (10 μg/cm^2^ surface area). The medium was replaced with PXGL on the following day and changed daily until dome-shaped colonies emerged (approximately 4–5 days after seeding). To enrich naïve-like human cells, SUSD2-positive cells with the labeled fluorescent reporter expression were sorted either with FACS or MACS as described above. The SUSD2-positive human cells were cultured on MEF feeder cells in PXGL. Phase contrast and fluorescence images in live cells were acquired using an all-in-one fluorescence microscope BZ-X800 (Keyence).

For the coculture experiments using human iPSCs lentivirally expressing shRNA (puromysin-resistant), day 5 coculture of human iPSCs with mouse ESCs (puromycin-sensitive) was passaged with Accutase on drug-resistant MEF feeder DR4 (Applied StemCell, Inc.) in PXGL containing Y-27632, Geltrex (10 μg/cm^2^ surface area), and 1.0 μg/ml puromycin. On the next day, the medium was replaced with PXGL supplemented with puromycin and changed every day for 7 days, which induced the negligible existence of mouse ESCs.

To track the mESC-derived mRNAs during coculture, *in vitro* transcribed, Cy5 labelled GFP mRNA (OZ Biosciences) was used. Transfection of the Cy5 GFP mRNA was performed with Lipofectamine MessengerMax (ThermoFisher Scientific) per manufacturer’s instruction. The next day the transfected mESCs were washed three times with PBS to use the coculture with H9 hESCs in 2i+LIF condition. L-778123 dihydrochloride (Tokyo Chemical Industry) was used in the indicated experiments.

### Transwell culture and conditioned medium assay

For transwell coculture experiments, Millicell cell culture 0.4 μm PET inserts (Millipore) were used by placing them into 12-well plates. Human PSCs were seeded on the bottom well coated with Matrigel or iMatrix-511, while mouse ESCs on the top insert coated with gelatin. The number of seeded cells used was same as in the direct coculture experiments. Conditioned medium was collected from day-3–5 cocultures and separate cultures, filtered through a 0.45 μm Minisart filter (Sartorius) to remove cell debris, and then used to culture human PSCs.

### Generation of fluorescent reporter lines

The plasmids used in this study are listed in **Supplementary Table 2**. To construct the donor plasmids for AAVS1-targeting knock-in in human PSCs, the sequences of fluorescent reporters were PCR-amplified, and then integrated into the multi-cloning site of PCR-amplified linearized pAAVS1-P-CAG-mCh (Addgene plasmid #80492), an AAVS1 targeting vector containing puromycin resistant gene, by using In-Fusion cloning kit (TaKaRa). For pAAVS1-Nst-MCS (Addgene #80487), an AAVS1 targeting vector containing neomycin resistant gene, the reporter sequences together with CAG promoter were PCR-amplified and integrated into the multi-cloning site of PCR-amplified linearized vector. To construct the donor plasmids for Rosa26-targeting knock-in in mouse ESCs, the left and right homology arms (∼1 kb) corresponding to mouse *Rosa26* locus were amplified from genomic DNA of J1 mESC line. Using pAAVS1-P-CAG as a backbone, these homology arms were swapped with the human AAVS1 homology arms, and the sequences of reporter cassettes were integrated by using In-Fusion cloning kit. PCR-amplified regions and In-Fusion junctions were verified by Sanger sequencing. pXAT2 (Addgene plasmid #80494) was used as the AAVS1 sgRNA expression CRISPR-Cas9 plasmid. The sgRNA sequence targeting mouse *Rosa26* is the following: 5’-gTGGGCGGGAGTCTTCTGGGC-3’. The designed sequence was cloned into the BbsI sites of pEF-SpCas9HiFi-2APuro-U6-gRNA, which is modified from the original pCS2-U6-gRNA-FN-Cas9-N-P2A-Puro(56) (kindly provided by S. Takahashi, The University of Tokyo, Japan) where the CMV promoter was swapped with EF1α and introduction of R691A mutation into SpCas9 for high-fidelity Cas9 protein(57). The sgRNA were verified by Sanger sequencing. For gene targeting, the donor plasmids (8 μg) and Cas9 and sgRNA expression plasmids (2 μg) were co-transfected by electroporation (NEPA21, Nepa Gene Co. Ltd) into 1 × 10^6^ cells, which were then divided and plated under feeder-free condition for 48 h before initiating drug selection (0.5 μg/ml puromycin, nacalai tesque; G418, Fujifilm Wako Pure Chemical Corp.). Eight to nine days after plating, the drug-resistant cells were pooled and passaged. Clones were isolated with CELL PICKER (SHIMADZU) or by manual picking, and transferred with gentle dissociation into a 96-well culture plate. Isolated clones were incubated for 6-10 days. At 80% confluency, the cells were detached with Accutase. Cell suspensions were split for freezing as well as genomic DNA isolation. These clones were genotyped by PCR and sequencing before final verification by Southern blotting.

### Flow cytometry

The fluorescent dye-conjugated antibodies (**Supplementary Table 3**) and Ghost Dye Violet 450 (Tonbo Biosciences) were mixed in Cell Staining Buffer (BioLegend), and applied to 100 μl of 2– 5×10^5^ of single cells per reaction. Cells were incubated for 30 min on ice in the dark followed by washing three times with Cell Staining Buffer. The stained cells were analyzed with a BD LSRFortessa cell analyzer (BD Biosciences) or a BD FACSAria III flow cytometer. Data was analyzed using FlowJo V10.8.0 software (FlowJo, LLC).

### Immunofluorescent microscopy

hiPSCs were plated on Matrigel- and MEF feeder-coated Lab-TekII chamber slide (Nunc) or Cell Desk LF1 (Sumitomo Bakelite) coated with Matrigel, iMatrx-511, or MEF feeder, and then cultured under 5% O_2_ at 37°C. The cells were fixed with 4% paraformaldehyde (nacalai tesque) for 10 min at room temperature followed by permeabilization with 0.2% Triton X-100 (Sigma Aldrich) in PBS for 5 min. Cells were then blocked with MAXblock blocking medium (Active Motif) or 3% BSA in PBS for 1 hr. Primary and secondary antibodies were diluted in MAXblock blocking medium, and were applied for 1 hr and 20min, respectively. DNA was counter-stained with 1 μg/ml DAPI (ThermoFisher Scientific) for 15 min. The samples were washed twice with PBS between each step. Images were taken with FV3000 confocal microscope (Olympus). Antibody details can be found in **Supplementary Table 3**.

To preserve and visualize the membrane protrusions, the different staining protocol was applied. Following the coculture, cell membrane was stained in live cells with CellBrite Fix 640 Membrane Labeling Kit (Biotium). The cells were then fixed in 2% paraformaldehyde and 0.1% glutaraldehyde (nacalai tesque) for 15 min at room temperature followed by permeabilization with 0.2% Triton X-100 (Sigma Aldrich) in PBS for 5 min. After fixation, cells were washed three times in PBS for 5 min each wash followed by incubation with 50 mM NH_4_Cl in PBS for 15 min. Cells were then permeabilized in 0.1% Triton X-100 in PBS for 5 min. After three times wash with PBS, cells were stained with Alexa Fluor-conjugated phalloidin (1:1,000) for 1 hr at room temperature. The samples were washed three times with PBS and mounted with PermaFluor Aqueous Mounting Medium (Lab Vision Corp.). Images were taken with SP8 confocal miscroscope (Leica) with HyD detectors being in photon counting mode.

### RNA extraction and RT-qPCR

Total RNA was extracted using RNeasy Mini Kit (QIAGEN) or FastGene RNA Basic Kit (NIPPON Genetics, Co., Ltd.). For FACS sorted samples, about 1×10^3^ – 1×10^4^ cells were subjected to total RNA isolation using NucleoSpin RNA Plus XS (MACHEREY-NAGEL GmbH & Co. KG). The first-strand cDNA was synthesized by SuperScript III (ThermoFisher Scientific) or ReverTra Ace qPCR RT Master Mix with gDNA Remover (TOYOBO Co., Ltd.). RT-qPCR was performed using THUNDERBIRD Next SYBR qPCR Mix (TOYOBO Co., Ltd.) with specific primer pairs listed in **Supplementary Table 4**. For quantification of human- or mouse-specific transcripts, RT-qPCR was performed using THUNDERBIRD Probe qPCR Mix (TOYOBO Co., Ltd.) with primer pairs and probes targeting for each gene listed in **Supplementary Table 4**. The expression levels were normalized against β-actin genes (*ACTB* in human; *Actb* in mouse). The data were collected from at least three independent biological replicates using StepOnePlus or QuantStudio 3 Real-Time PCR Systems (ThermoFisher Scientific).

### Lentiviral shRNA knockdown

To design shRNAs targeting mouse *Tfcp2l1*, *Kl4*, and *Tfap2c* genes, but not human ones, both human and mouse transcripts were aligned by using Clustal W(58). Among the less conserved regions, the shRNA candidate sequences were designed by using GPP Web Portal provided by the Broad Institute (https://portals.broadinstitute.org/gpp/public/seq/search). The shRNA sequences were cloned into the AgeI-EcoRI site of pLKO.1. The oligonucleotides used for the cloning are listed in **Supplementary Table 4**. The lentiviral packaging was produced in HEK293T cells transfected with pCAG-HIVgp and pCMV-VSV-G-RSV-Rev packaging plasmids (provided by RIKEN BRC DNA Bank) via TransIT-2020 (TaKaRa). In order to conduct lentiviral infection, the virus-containing medium supplemented with 8 μg/ml of polybrene (InVivogen) was added into human iPSC culture, and the culture plates were supn at 1,000 × *g* for 30 min at 32℃. After the incubation for 24 hr, the medium was replaced followed by the selection with 1 μg/ml of puromycin.

### RNAscope

Cells were plated on Cell Desk LF1 (Sumitomo Bakelite) coated with gelatin, iMatrx-511, or MEF feeder cells. The cells were fixed with 4% paraformaldehyde in phosphate buffer for 10 min at room temperature. RNAscope was carried out using RNAscope Fluorescent Multiplex Reagent kit v1 (Advanced Cell Diagnostics, Inc.) according to the manufacture’s protocol. RNAscope probes used are listed in **Supplementary Table 4**. For counter-staining with antibodies, the coverslips from RNAscope were further blocked with BSA blocking buffer (3% BSA, 0.1% Triton X-100 in PBS) for 1 hr at room temperature. After incubation with primary antibodies diluted in BSA blocking buffer at 4°C over night, they were washed with PBS three times followed by incubation with Alexa Fluor-conjugated secondary antibodies and Hoechst 33342 diluted in BSA blocking buffer at room temperature for 1 hr. After washing with PBS three times, the samples were mounted with ProLong Diamond Antifade Mountant (Invitrogen). Images were taken with SP8 confocal microscope (Leica) or FV3000 confocal microscope (Olympus).

Although fine structures of intercellular protrusions were well preserved in the presence of low concentration of glutaraldehyde as a fixative, the staining condition prevented the detection of mRNAs by RNAscope as discussed previously(59). However, in the standard staining condition described above, which less retained the protrusions during the processing, we could detect the mRNA spots in the remaining protrusions as shown in **Fig. 3B,C**.

### Image processing and quantification

Image processing and quantification was carried out using Fiji software.

### RNA-sequencing

Total RNA was purified using an RNeasy Micro Kit (Qiagen) according to the manufacturer’s instructions. RNA quality and quantity were checked using Bioanalyzer (Agilent) and Qubit (Life Technologies) machines, respectively. The initial amplification step was performed with the NuGEN Ovation RNA-Seq System v2, which facilitates an assay that amplifies RNA samples and creates double-stranded cDNA. Libraries were then created with the Nextera XT DNA Sample Preparation Kit (Illumina) and sequenced with an Illumina HiSeq 2500 system. RNA-seq data analyses were performed using the BioWardrobe Experiment Management System [https://github.com/Barski-lab/biowardrobe]. Briefly, reads were mapped to the mm10 genome using TopHat [version 2.0.9] and assigned to RefSeq genes (which have one annotation per gene) using the BioWardrobe algorithm. PCA was performed using the top 1000 most variable genes across experimental condition. The first and second principal components were plotted. Differential gene expression analyses were performed via DESeq2 in the BioWardrobe environment. For gene ontology analyses, the Database for Annotation, Visualization and Integrated Discovery (DAVID) was used.

To separate mouse and human sequences, the function bbsplit from the bbtools suite (BBTools) was used to bin each read from the RNA-seq fastq into a separate fastq files based on whether the read was specific to the human (hg19) or mouse (mm10) genome. To create the most stringent conditions, perfect mode was set to true and all ambiguous reads were discarded. Reads that were specific to mouse within the human cells were aligned to the mm10 genome using STAR (v2.5.1b). The resulting aligned sam file was sorted and converted to a bam file using the samtools sort function. Reads for each gene were counted using the R function summarizeOverlaps from the GenomicAlignments library using the counting mode of Union. DESeq2^2^ was used to quantify differential gene expression among samples.

To compare gene expression profile of human naïve cell lines established in this study with originally reported naïve PSC lines, deposited sequencing data of Shef6-primed (accession numbers: ERR1924246, ERR1924247, ERR1924248.) and Shef6-cR (accession numbers: ERR1924234, ERR1924235, ERR1924236.) were obtained from the European Nucleotide Archive. The all sequencing data were processed to remove adaptor sequences with Trimmomatic(60). Mapping the sequences to the human genome (GRCh38) was performed with STAR(61). Transcript abundances were quantified and converted to count data using RSEM(61). After the low count genes were excluded, raw counts were normalized using the variance stabilizing transformation and differentially expressed genes were identified using the DEseq function in the DESeq2 R package(62). Non-biological batch effects related to the experimental setting differences was compensated using the ComBat function as implemented in the sva R package(63). The gene ontology analysis and over-representation analysis combined with KEGG pathway were performed using ShinyGO(64) and NetworkAnalyst(65), respectively.

To compare the transcriptome in hESCs over mESC coculture-mediated conversion, we prepared libraries from total RNAs with NEBNext Poly(A) mRNA Magnetic Isolation Module (New England Biolabs) and NEBNext Ultra TMII Directional RNA Library Prep Kit (New England Biolabs). Libraries were sequenced with an Illumina NovaSeq 6000 to obtain 150 bp ×2 paired-end reads. The quality of the raw paired-end sequence reads was assessed with FastQC (v0.11.7). Low quality bases and adapter sequences were trimmed by Trimmomatic software (v0.38). The trimmed reads were aligned to both the human hg38 and mouse GRCm38 reference genomes using STAR aligner (v2.7.4a) in two-pass mode, and mouse contamination reads were filtered from the human alighed reads using the XenofilteR software (v1.6). The filtered bam files were used to estimate the abundance of uniquely mapped reads with featureCounts (v1.6.3)(66). To extract the differentially expressed genes along the mESC coculture-mediated conversion, we performed likelihood ratio test for the samples by using DESeq2 package with adjusted *P* values <0.01 and log2 fold change >=2. To examine the expression patterns across sample groups, we used the degPatterns function from the DEGreport R package (v3.19). The gene ontology analysis was performed using the clusterProfiler R package(67).

### RNA-seq variant calling and biallelic expression analysis of imprinted genes

To analyze the loss of imprinting we identified biallelic expression of imprinted genes through calculation of their allelic ratios and biallelic scores from the bulk RNA-seq datasets of PB004 human PSCs as described previously(33) with minor modifications. Briefly, the fastq files were trimmed by Trimmomatic software (v0.38) followed by alignment in parallel to both the human GRCh38 and mouse GRCm38 reference genomes using STAR (v2.7.4a). Mouse contamination reads were filtered from the human aligned reads using the XenofilteR (v1.6)(66). The filtered bam files were subjected to two parallel procedures: gene count matrix generation with featureCounts (v1.6.3) and variant calling. For the latter, the duplicated reads were filtered by Picard MarkDuplicates (https://broadinstitute.github.io/picard/) followed by the extraction of variants with small polymorphisms (SNPs, small indels, MNPs, and composite events) through the GATK software (SplitNCiger, BaseRecalibrator and ApplyBSQR). The identified small polymorphisms were filtered with the depth >=10 in at least one sample as well as annotated using SnpEff(68). To determine the loss of imprinting for imprinted genes, we first identified the SNVs that were along exons of isoforms with IDs of dbSNP. SNVs with no annotated ID and coverage of less than 8 read coverages were filtered out. A gene was considered to lose its imprinting if at least one SNV along its exons had a minimum biallelic ratio value of 0.2 (the number of counts for the allele with the lower coverage divided by the number of counts for the allele with the higher coverage).

## Accession Numbers

RNA seq data hve been deposited under accession number GSE216348 and GSE216338 in the Gene Expression Omnibus (GE) of NCBI.

