## Supplementary Figures and Legends for "Inter-cellular mRNA Transfer Alters Human Pluripotent Stem Cell State"

### Supplementary Figure Legends

**Fig. S1. Mouse ESC co-culture potentiates adaptation of human PSCs in 2i+LIF condition.**

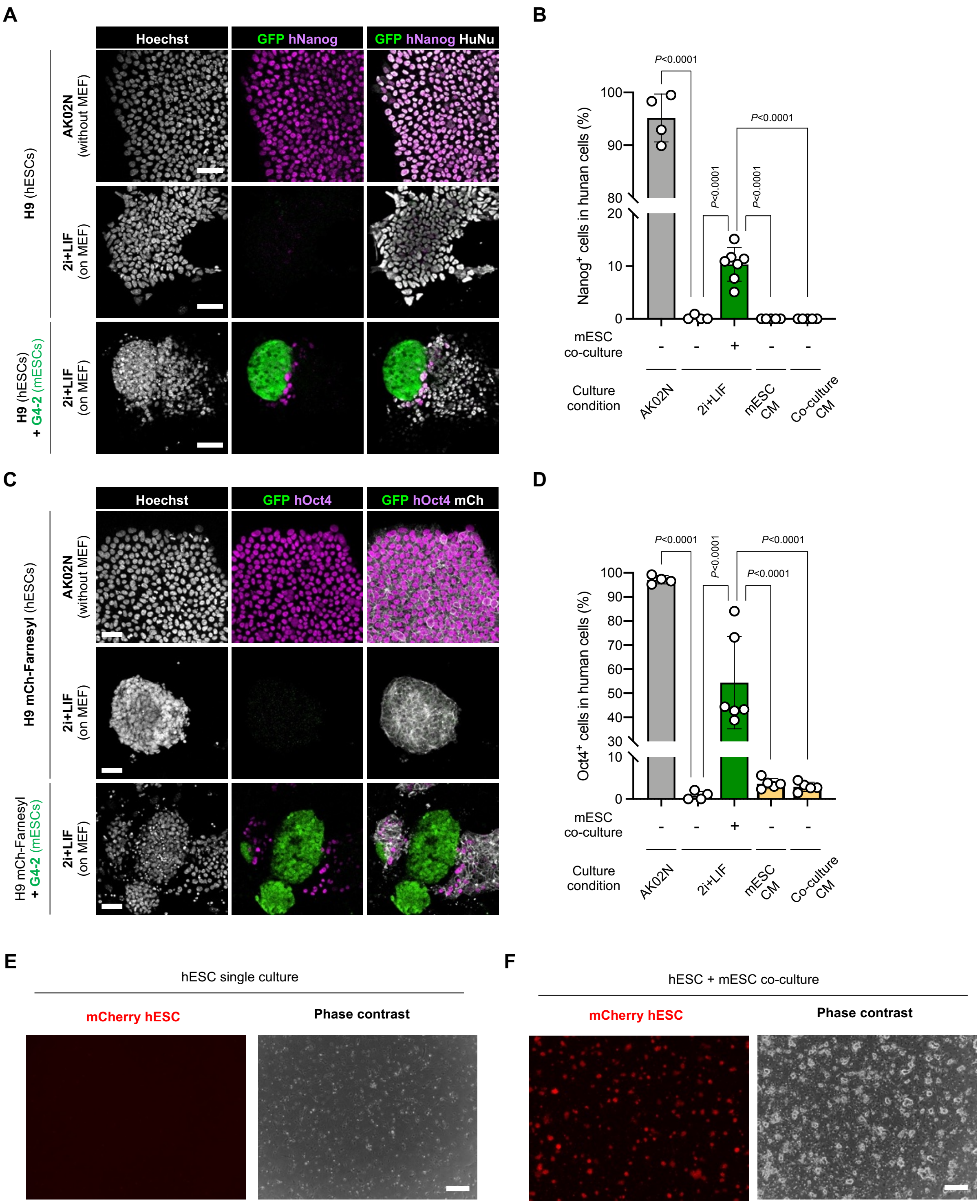


**Figure S1. Mouse ESC co-culture potentiates adaptation of human PSCs in 2i+LIF condition.**
(**A**) Representative immunofluorescence images of day-5 separately cultured H9 hESCs in primed culture medium (AK02N) or 2i+LIF, and cocultured H9 cells and G4-2 mESCs (green) in 2i+LIF. Cells were stained with antibodies against human-specific nuclear antigen (HuNu) and human-specific Nanog (hNanog). Scale bar, 50 μm.

(**B**) Quantification of hNanog-positive H9 hESCs in (**a**) (n ≧ 4 randomly selected fields). For separately cultured conditions without mESCs, conditioned medium (CM) collected from mESC separate culture in 2i+LIF or co-culture of H9 and G4-2 cells in 2i+LIF was also used. The data are representative of three independent experiments. Average ± SD is shown.
(**C**) Representative immunofluorescence images of day-5 separately cultured H9 mCherry-Farnesyl hESCs in primed culture medium (AK02N) or 2i+LIF, and cocultured H9 cells and G4-2 mESCs (green) in 2i+LIF. Cells were stained with antibodies against human-specific nuclear antigen (HuNu) and human-specific Oct4 (hOct4). Scale bar, 50 μm.
(**D**) Quantification of hOct4-positive H9 hESCs in (**c**) (n ≧ 4 randomly selected fields). For separately cultured conditions without mESCs, conditioned medium (CM) collected from mESC separate culture in 2i+LIF or coculture of H9 and G4-2 cells in 2i+LIF was also used. The data are representative of three independent experiments. Average ± SD is shown.
(**E**) Representative images of hESCs (H9 mCherry-Farnesyl) without mESC coculture captured at the indicated time points during the culture in 2i+LIF and PXGL. There remained few growing cells at second passage after human cell sorting of the 2i+LIF culture. Scale bar, 100 μm.
(**F**) Representative images of naïve-converted hESCs via mESC coculture. The converted cells continued to grow in PXGL, and the image shown is the culture at passage 2 post SUSD2 FACS sorting. Scale bar, 100 μm.

### Fig. S2. Intercellular mRNA transfer between human and mouse PSCs.

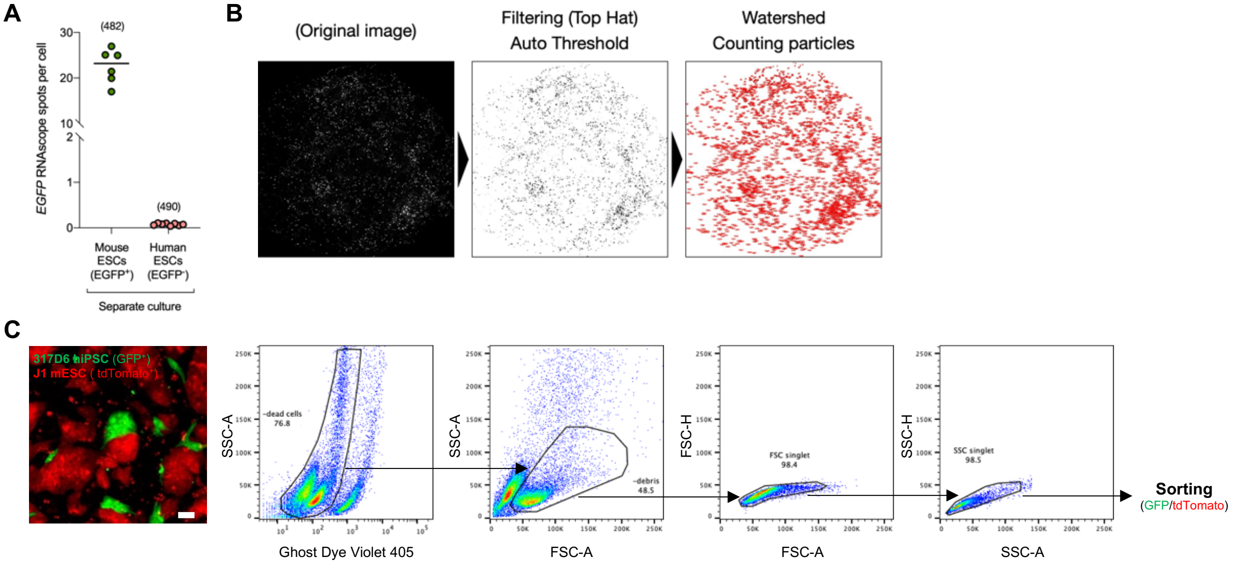


**D**

| Sample | % human | % Mouse | % cells outside human and mouse gates |
|---|---|---|---|
| Human cells sorted from mix with co-culture | 91.8 ± 0.72 | 0.00 ± 0.00 | 8.17 ± 0.72 |
| Mouse cells sorted from mix with co-culture | 0.00 ± 0.00 | 99.9 ± 0.00 | 0.10 ± 0.00 |
| Human cells sorted from mix without co-culture | 99.6 ± 0.00 | 0.00 ± 0.00 | 0.40 ± 0.00 |
| Mouse cells sorted from mix without co-culture | 0.00 ± 0.00 | 98.8 ± 0.06 | 1.23 ± 0.06 |

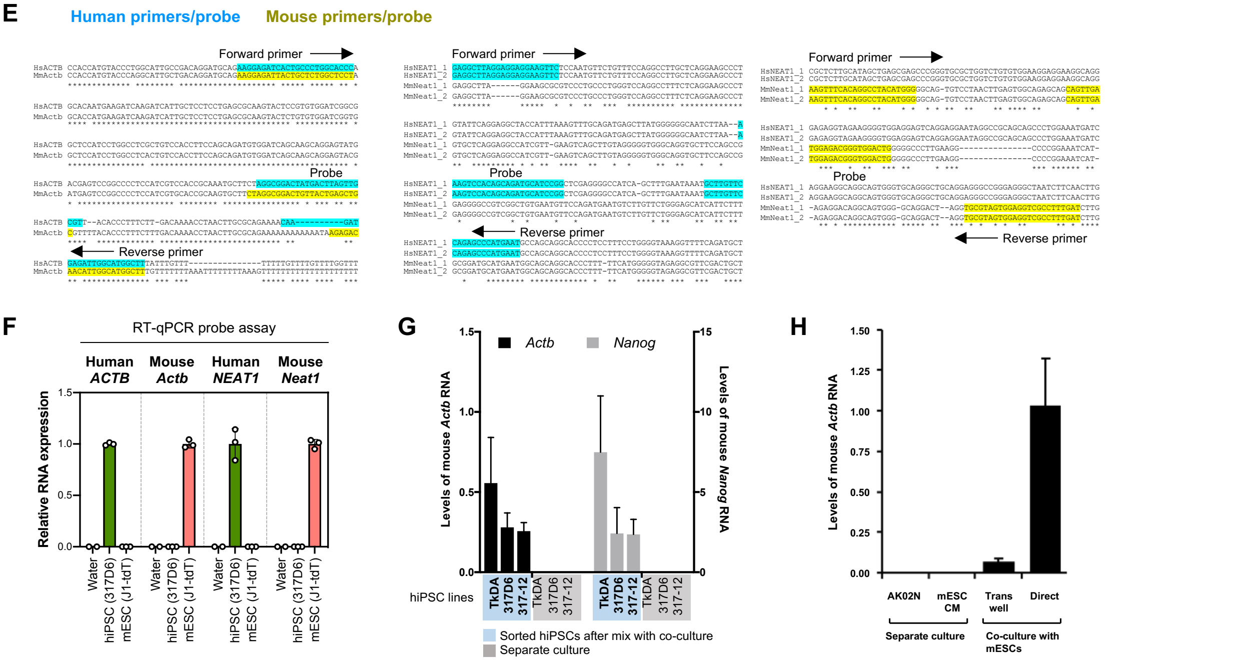


**Supplementary Figure 2. Intercellular mRNA transfer between human and mouse PSCs.**
(**A**) Quantification of the number of *EGFP* RNA spots in the separately cultured mESCs (EGFP-positive) and hESCs (EGFP-negative). The randomly selected fields (n=6-8) of each sample were analyzed, and the RNAscope spots in cells were counted (the number of cells analyzed is indicated in parentheses of the graphs). The black horizontal lines indicate the mean of the distribution.

(**B**) Spot detection and counting of RNAscope signals. Shown are the image processing in Fiji where filtering (Top Hat) and auto threshold were used to generate the binary image and watershed and counting particles functions were to count the RNAscope spots with the size pixel filtering.
(**C**) Representative FACS analysis to isolate live, singlet cells from hiPSC and mESC coculture. Coculture at day 5 of GFP-positive 317D6 hiPSCs and tdTomato-positive J1-tdT mESCs (left image, bar, 200 μm) was dissociated and stained with a dead cell dye (Ghost Dye Violet 405), followed by gating live, FSC/SSC-singlet cells that were further sorted into $GFP^+$ $tdTomato^-$ or $GFP^-$ $tdTomato^+$ cells.
(**D**) Percentage of separated human or mouse cells after FACS described in **A**. The sorted cells were analyzed by using the same gating as cell sorting. The data from three independent experiments are represented as the average ± SD.
(**E,F**) Design of primers and probes for RT-qPCR analysis to detect β-actin mRNA (*ACTB* in human, *Actb* in mouse) and NEAT1 (*NEAT1* in human, *Neat1* in mouse) long non-coding RNA in a human (blue)- or mouse (yellow)-specific manner (**E**). RT-qPCR either of hiPSC or mESC alone with the designed primers and probes validating the human/mouse-specific detection (**F**). Average ± SD (n=3) is shown.
(**G**) RT-qPCR analysis of mouse-specific *Actb* (black bars) and *Nanog* (grey bars) expression levels in the sorted hiPSCs after the co-culture with mESCs for 5 days. The three hiPSC lines, TkDA, 317D6, and 317-12, were used in this experiment. The values indicate the relative detected levels in hiPSCs either derived from sorted fractions after the mix with coculture or from separate culture. Average ± SD (n=3) is shown.
(**H**) RT-qPCR analysis of mouse-specific *Actb* expression levels in the sorted hiPSCs from direct or transwell coculture with mESCs and in the separate culture in primed or mESC conditioned medium (CM) condition. Average ± SD (n=3) is shown.

### Fig. S3. Functional characterization of mouse ESC-derived mRNA in human iPSC

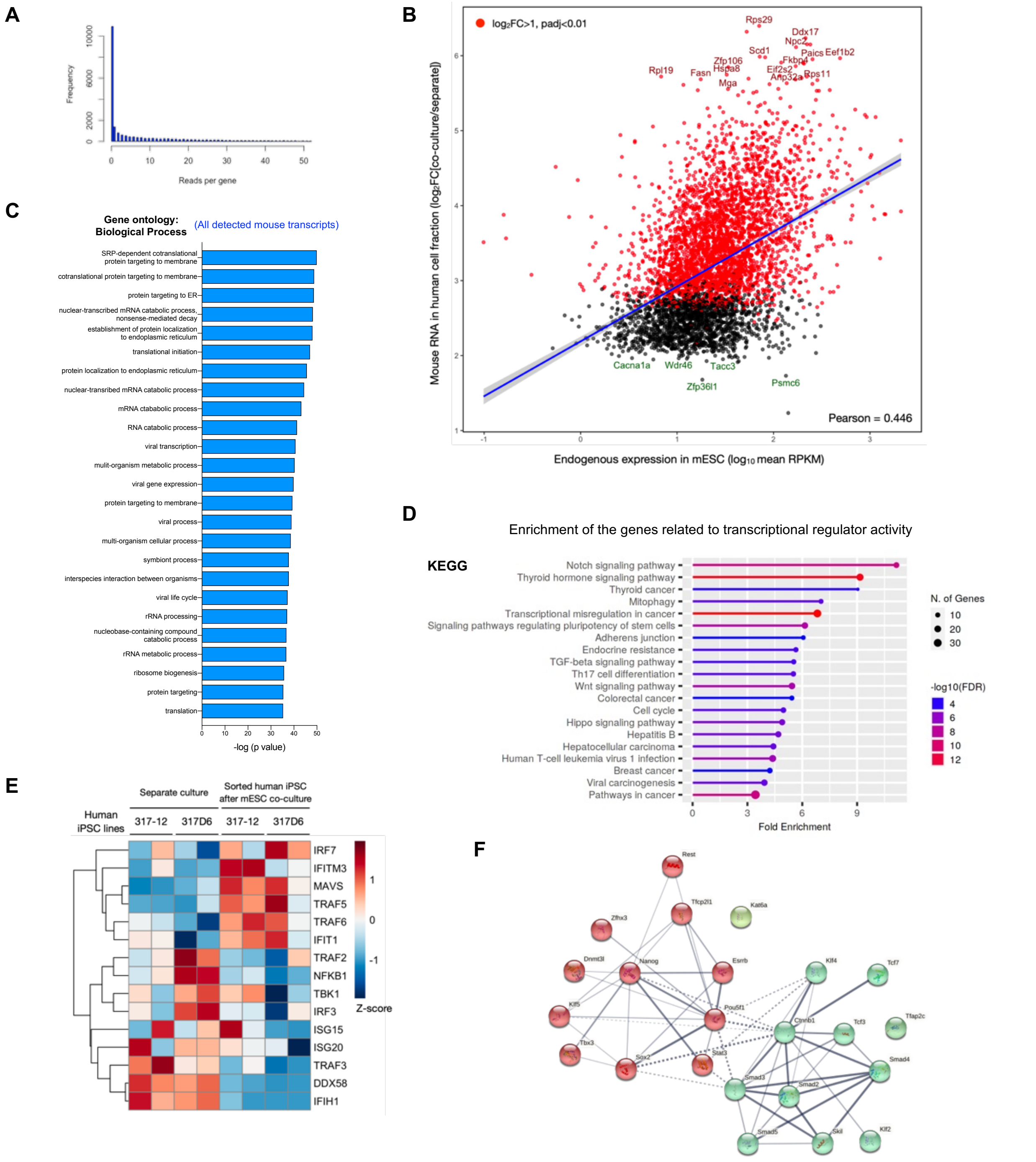


**Supplementary Figure 3. Functional characterization of mouse ESC-derived transferome in human iPSCs.**
(**A**) Read distribution of mouse-specific genes detected in the sorted hiPSCs after co-culture with mESCs.
(**B**) Correlation between the efficiency of mouse RNA transfer to hiPSCs and the endogenous level of gene expression in donor mESCs. The mouse genes upregulated in mESC coculture condition identified in **Fig. 2A** were subjected to correlation and linear regression analysis, where the transferred mouse RNA ratio (log2 fold change of coculture over separate condition) was compared with endogenous expression in separate cultured mESCs per a mouse gene. The differentially expressed genes with log2 fold change >1 and adjusted *P* value <0.01 are labelled in red. The blue line indicates the linear regression line with the upper and lower bounds of 95% confidence internal. Pearson coefficient is also indicated.
(**C**) Gene ontology (GO) analysis of the mouse genes enriched in the sorted hiPSCs after coculture with mESCs. The list of differentially expressed genes in RNA-seq (Figure 3A) was analyzed by DAVID (database for annotation, visualization, and integrated discovery) for the GO annotation. The top 25 terms for the term family of Biological Processes are shown along with their -log (*p* value).
(**D**) The top-ranked KEGG pathways that are overrepresented within the set of mouse transcription regulator activity genes enriched in the sorted hiPSCs after the coculture with mESCs. The fold enrichment with significantly enriched genes (FDR<0.05) was computed. The color

chart shows the FDR values for enrichment for the respective pathways (-$\log_{10}$(FDR)), while the size of dots corresponds to the number of genes assigned to each pathway.
(**E**) Expression heatmap of human genes related to the RIG-1 pathway in hiPSCs sorted from mESC coculture or separate culture condition.
(**F**) Protein-protein interaction network of transcription factors grouped in the KEGG pathway regulating pluripotency of stem cells enriched in the mouse genes of hiPSCs after co-culture with mESCs. Interactions with a high confidence (>0.7) are visualized by STRING. The transcription factors are clustered using the MCL algorithm, each color represents a cluster.

### Fig. S4. Intercellular transfer of transcription factor-coding mRNAs from mouse ESCs into human PSCs.

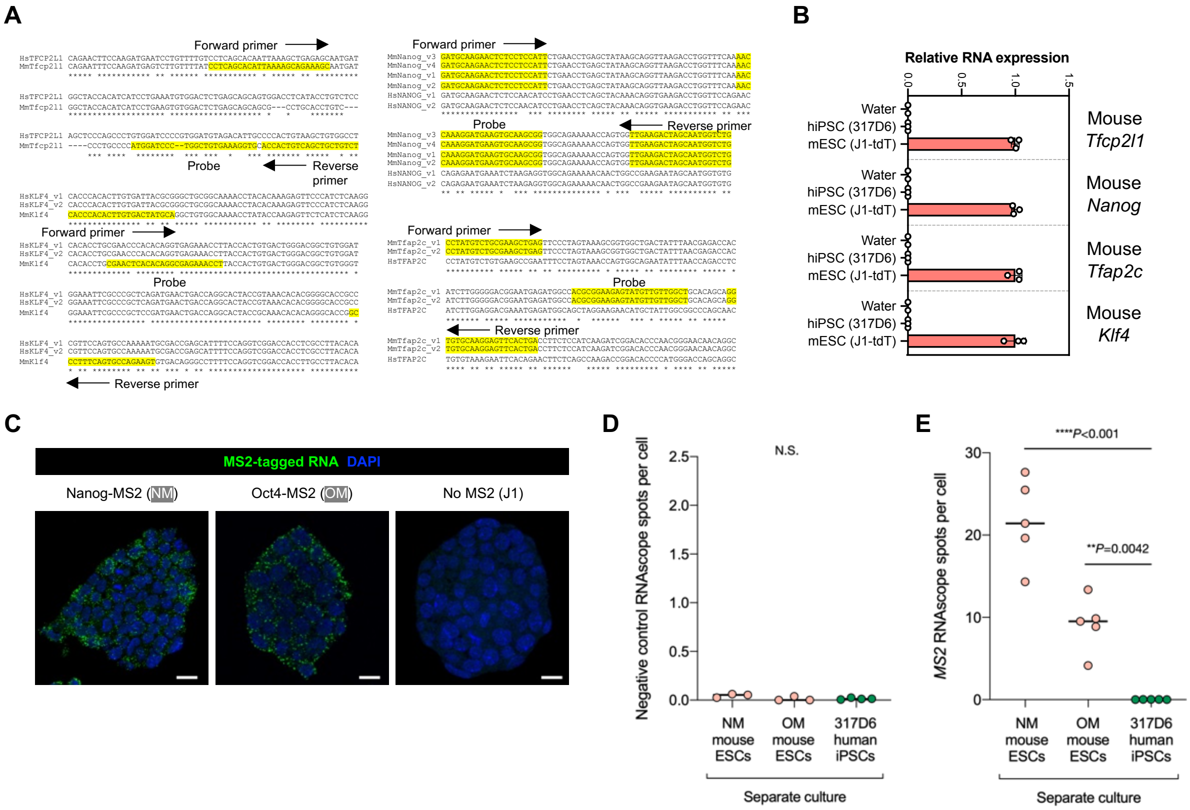


**Supplementary Figure 4. Intercellular transfer of transcription factor-coding mRNAs from mouse ESCs into human iPSCs.**
(**A**) Design of primers and probes for RT-qPCR analysis to detect mouse *Tfcp2l1*, *Klf4*, *Nanog*, and *Tfap2c* mRNAs in a mouse (yellow)-specific manner.
(**B**) RT-qPCR detection of mouse *Tfcp2l1*, *Klf4*, *Nanog*, and *Tfap2c* transcripts specific for mouse by using the corresponding primers and probes. Total RNAs extracted from hiPSCs and mESCs were tested in this experiment. Average ± SD (n=3) is shown.
(**C**) Representative RNAscope images of MS2-tagged RNA in mESCs expressing Nanog-MS2 (NM) and Oct4-MS2 (OM). The J1 cell line without MS2 reporter was set as a negative control. The cells were also counter-stained with DAPI. Scale bar, 10 μm.
(**D**) Quantification of the number of RNA spots stained with negative control probe in the mESCs (NM and OM) and hiPSCs (317D6). The randomly selected fields (n=3-4) of each sample were analyzed, and the RNAscope spots in cells were counted. The black horizontal lines indicate the mean of the distribution.
(**E**) Quantification of the number of MS2-tagged RNA spots in the mESCs (NM and OM) and hiPSCs (317D6). The randomly selected fields (n=5) of each sample were analyzed, and the RNAscope spots in cells were counted. The black horizontal lines indicate the mean of the distribution.

### Fig. S5. Effect of L-778123 on intercellular membrane protrusion and mESC-derived RNA transfer

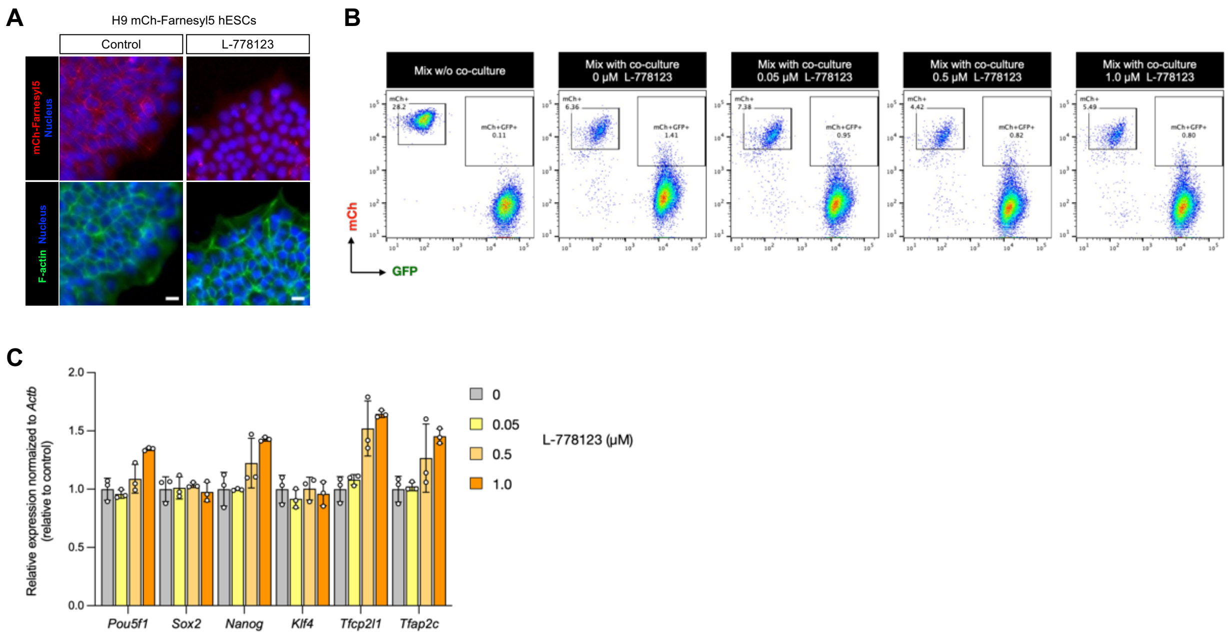


**Supplementary Figure 5. Effect of L-778123 on intercellular membrane protrusion and mESC-derived RNA transfer.**
(**A**) Localization of the mCherry-Farnesyl5 reporter expressed in H9 hESCs in response to L778123. H9 mCh-Farnesyl5 cells were cultured in primed condition in the presence or absence of L-778123 (0.5 μM). The cells were stained with phalloidin and Hoechst 33342 to visualize F-actin and nuclei, respectively. Scale bar, 10 μm.
(**B**) Representative FACS analysis to isolate mCherry-positive human cells from the coculture with mESCs treated with L-778123 at the indicated concentrations in **Fig. 5G**. The panels shown are from cells gated as live, FSC/SSC-singlet cells.
(**C**) RT-qPCR analysis of *Pou5f1*, *Sox2*, *Nanog*, *Klf4*, *Tfcp2l1*, and *Tfap2c* genes expressed in mESCs treated with L-778123 at the indicated concentrations. Relative values normalized with endogenous *Actb* expression are shown as an average ± SD (n=3).

### Fig. S6. Presence of mouse specific proteins inside human cells after mESC-derived mRNAs transfer.

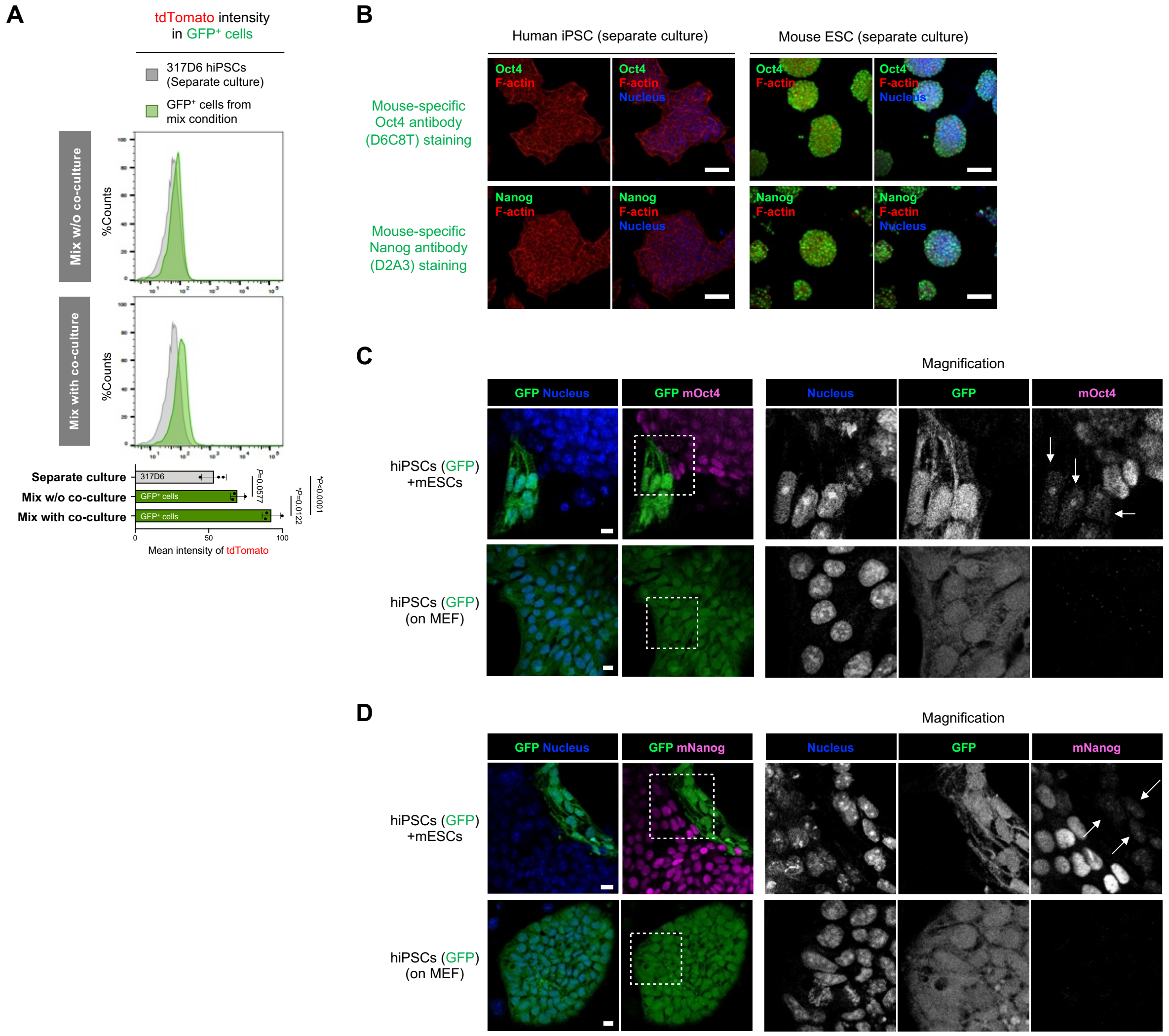


**Supplementary Figure 6. Presence of mouse specific proteins inside human cells after mESC-derived mRNAs transfer.**
(**A**) Quantification of tdTomato intensity in GFP-positive human iPSCs or GFP intensity in tdTomato-positive mouse ESCs in both mix with and w/o coculture conditions shown in **Fig. 1D**. The relative intensities of GFP or tdTomato compared with separate culture of human iPSCs or mouse ESCs are also shown as bar graphs (average ± SD, from three independent experiments).
(**B**) Immunofluorescence of human iPSCs and mouse ESCs with mouse-specific Oct4 and Nanog antibodies. Cells were counterstained with Hoechst 33342 and phalloidin to visualize nuclei and F-actin, respectively. Scale bar, 100 μm.
(**C,D**) Immunofluorescence of human iPSCs (GFP-expressing) coculture with mouse EScs or human iPSCs alone (on MEF) with mouse-specific Oct4 (**C**) and Nanog (**D**) antibodies, respectively. Magnification images highlight the presence of dim signals of mouse Oct4 and Nanog proteins inside human cells upon co-culture (white arrows). Scale bar, 10 μm.

**Fig. S7. Gene expression changes in human PSCs over the naïve-like conversion mediated by mouse ESC co-culture.**

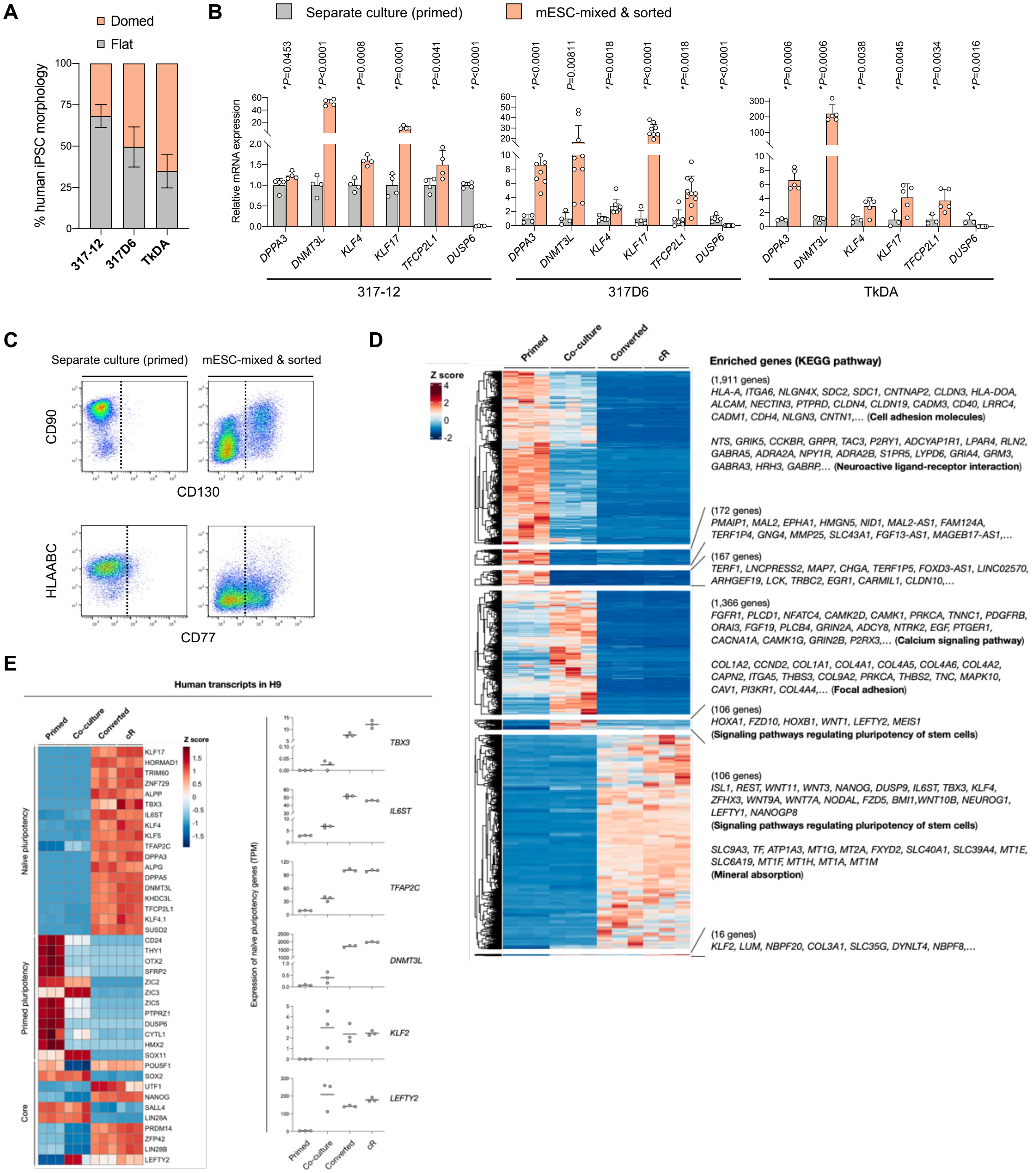


**Supplementary Figure 7. Gene expression changes in human PSCs over the naïve-like conversion mediated by mouse ESC co-culture.**

(**A**) Ratio of flat and dome-shaped colonies observed in three human iPSC lines (317-12, 317D6, TkDA) following the co-culture with mouse ESCs for 5 days. The data from three independent experiments are represented as the percentage of the observed colonies (average ± SD).
(**B**) RT-qPCR analyses of primed (*DUSP6*) and naïve (*DPPA3*, *DNMT3L*, *TFCP2L1*, *KLF4*, and *KLF17*) marker genes in three human iPSC lines (317-12, 317D6, TkDA) that were separately maintained in the primed state (separate culture), or were enriched by FACS after the co-culture with mouse ESCs for 5 days (n=3–10, average ± SD). The *P* values of each gene comparison are shown.
(**C**) FACS profiles of primed (CD99 and HLAABC) and naïve (CD130 and CD77) cell surface markers in the separately cultured, primed human iPSCs and the putative naïve-like human iPSCs expanded from the co-culture with mouse ESCs.
(**D**) RNA-seq analysis of human transcripts expressed in H9 mCh-Farnesyl5 hESCs upon G4-2 mESCs coculture-mediated naïve-like conversion. The differentially expressed genes are identified along isolated human cells from mESC coculture (“co-culture”) and subsequent

culture of emerging SUSD2⁺ cells as well as primed and chemical reset ("cR") cells. Seven clusters revealed by pattern clustering are shown, some of which are indicated by the consisting genes and the representative KEGG pathways from gene ontology analysis.
(**E**) Expression heatmap of selected core, primed and naïve pluripotency genes in H9 hESC transcriptome in (**D**) (left). The right graphs show the expression (TPM) of pluripotency-related genes which are increased along the mESC coculture-mediated conversion. The black horizontal lines indicate the mean of the distribution (n=3).

### Fig. S8. Co-culture with mouse ESCs enables naïve-like conversion of human PSCs.

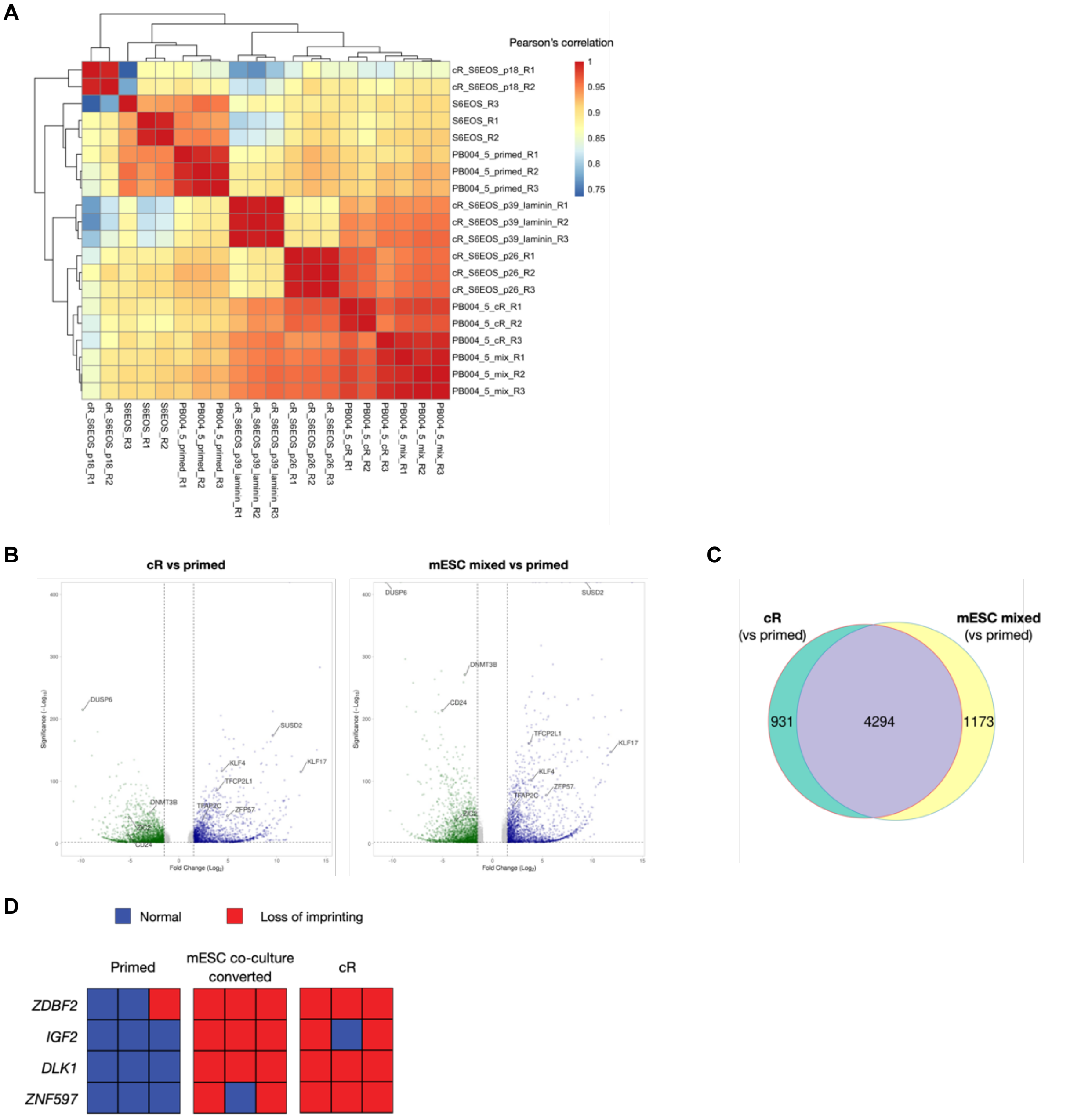


**Supplementary Figure 8. Co-culture with mouse ESCs enables naïve-like conversion of human PSCs.**
(**A**) Pearson correlation matrix of global gene expression in RNA-seq obtained from hiPSCs including primed cells (parental), cR cells, and converted cells via mESC co-culture and subsequent expansion, as well as previously published hESCs in primed and cR states (both on-feeder and feeder-free)[1].
(**B**) Volcano plots of RNA-seq to compare primed and cR hiPSCs (left), or primed and converted cells via mESC co-culture (right). Differentially expressed genes ([Fold Change] > 2, *p*-adj <0.01) are marked either in green or blue, which represents downregulated or upregulated genes, respectively. The genes related to naïve or primed pluripotency are highlighted.
(**C**) Venn diagrams show the overlap between differentially expressed genes in RNA-seq that are identified in cR and converted hiPSCs via mESC co-culture compared with primed state.
(**D**) Heatmap of biallelic ratio value of genes whose expression are observed to be biallelic in naïve/naïve-like condition of RNA-seq data in **Fig. 4F,G**.

**Fig. S9. Co-culture with rat naïve iPSCs induces intercellular mRNA transfer and subsequent naïve-like conversion of human PSCs.**

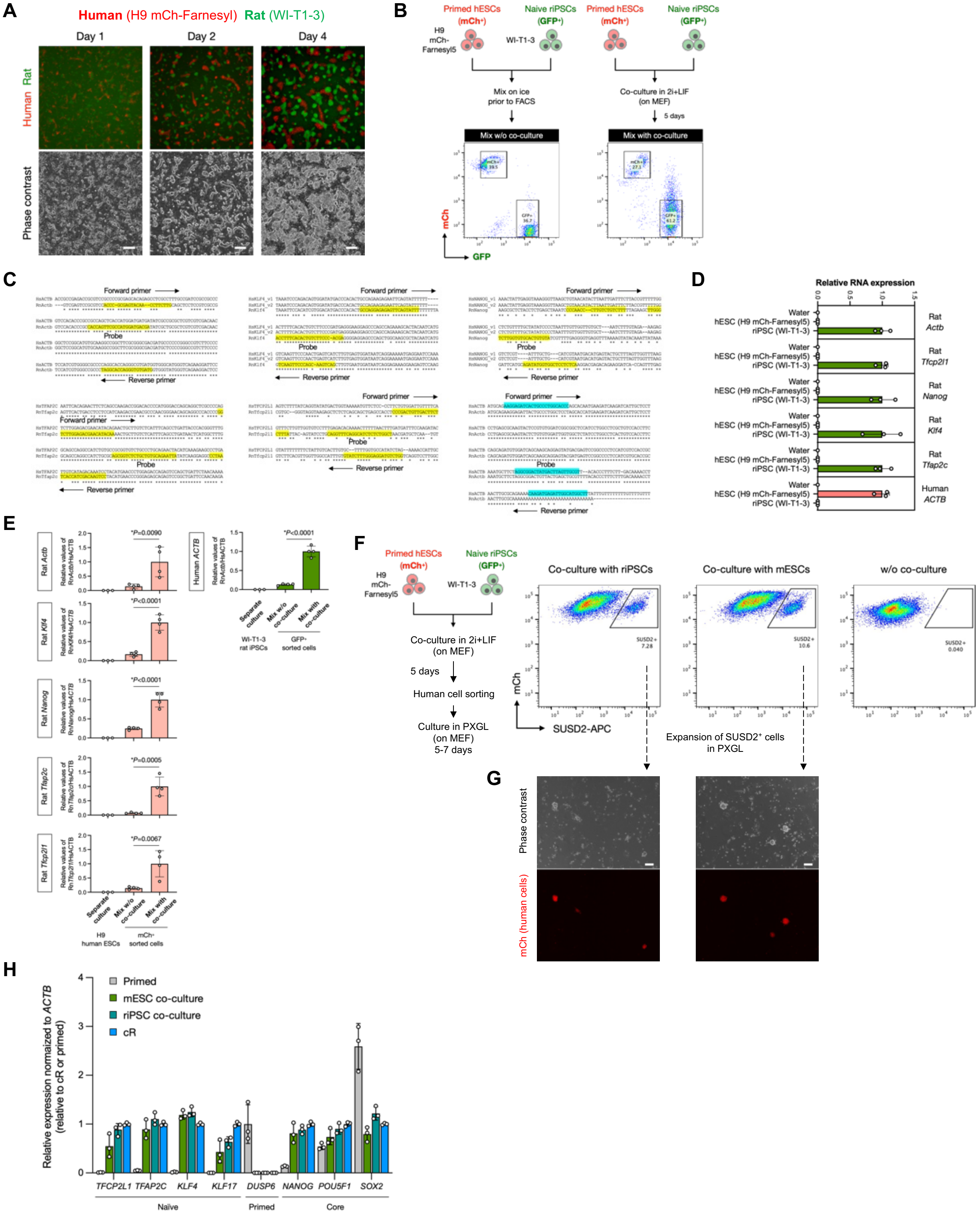

**Supplementary Figure 9. Co-culture with rat naïve iPSCs induces intercellular mRNA transfer and subsequent naïve-like conversion of human PSCs.**
(**A**) Representative images captured at the indicated time points during the coculture of primed hPSCs (H9 mCh-Farnesyl5, red) and naïve rat iPSC (WI-T1-3 riPSC, green). Scale bar, 200 μm.

(**B**) Experimental design for primed hESCs (mCherry-positive H9) and naïve riESCs (GFP-positive WI-T-3) co-culture followed by FACS. Representative FACS analyses of mixed cells were shown either from day-5 coculture (mix with coculture) or from mix on ice prior to FACS (mix w/o coculture). GFP$^{+}$/ mCherry$^{-}$ and GFP$^{-}$/ mCherry$^{+}$ cells were sorted for the subsequent RT-qPCR analyses.

### Fig. S10. Mouse transcription factor-specific shRNA expression did not affect naïve conversion of human iPSCs.

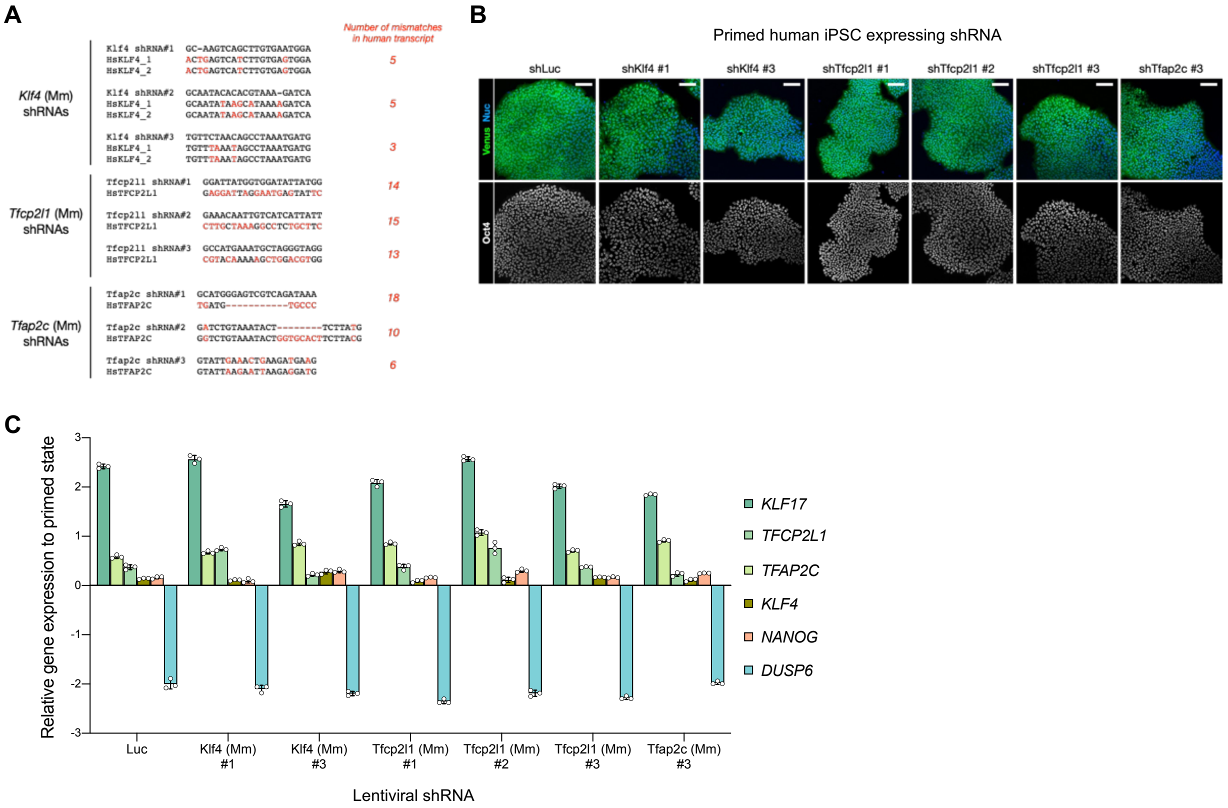


**Supplementary Figure 10. Mouse transcription factor-specific shRNA expression did not affect potency of naïve conversion of human iPSCs.**
(**a**) Design of shRNA sequences targeting mouse *Klf4*, *Tfcp2l1*, and *Tfap2c*. The target regions of the shRNAs in mouse transcripts were aligned to those corresponding to human transcripts. The mismatch bases found are in red, and the number of mismatches in human transcript is shown.
(**b**) Immunostaining of Oct4 in the human primed iPSCs (Venus-positive) in which lentiviral shRNAs targeting each mouse transcription factor are stably expressed. Scale bar, 100 μm.
(**c**) RT-qPCR analysis of naïve markers *KLF17*, *TFCP2L1*, *TFAP2C*, *KLF4*, and *NANOG*, and primed marker *DUSP6* in the chemically naïve-converted human iPSCs (cR) in which lentiviral shRNAs targeting each mouse transcription factor are stably expressed. The relative expression levels to their corresponding primed states are shown as Average ± SD (n=3).

**Supplementary Table Legends**

**Supplementary Table1.** Mouse transcription factor genes detected in sorted human iPSCs after mouse ESC co-culture by RNA-seq.

**Supplementary Table2.** Plasmids used in this study.

**Supplementary Table3.** Antibodies used in this study.

**Supplementary Table4.** Primers and probes for RT-qPCR, primers for shRNA, and probes for RNAscope used in this study.
